# Manipulating syntax without taxing working memory: MEG correlates of syntactic dependencies in a Verb-Second language

**DOI:** 10.1101/2024.02.20.581245

**Authors:** Simone Krogh, Liina Pylkkänen

## Abstract

The neural basis of syntax is notoriously difficult to study without working memory and lexico-semantic confounds. To tackle these challenges, we presented syntactic dependencies in Danish two-word sentences using Rapid Parallel Visual Presentation (RPVP), eliminating the temporal delay between displaced elements and their base-generated positions. Our stimuli involved various dependencies as a function of syntactic frame (declarative, yes/no question) and verb argument structure (unergative, unaccusative, alternating unaccusative). Behaviour was facilitated and magnetoencephalography responses were increased for sentences compared to list-controls in left fronto-temporal regions (231–407ms; 506–622ms), largely replicating the Sentence Superiority Effect of prior RPVP studies. Each cluster showed sensitivity to syntactic frame, with the earlier one likely reflecting basic structure detection and the second syntactic movement. No effects of verb argument structure were obtained. The RPVP paradigm’s sensitivity to both the sentence-list contrast and to certain grammatical details highlights its value for probing the neurobiology of syntax.

## INTRODUCTION

A core assumption of linguistic theories is that language processing is a tandem operation between syntax—the building and parsing of hierarchical structures—and the lexicon. Although large literatures have addressed the neural bases of both syntactic processing and lexical access, reaching agreement on higher levels beyond that of lexical access has proven challenging. Beginning with the by-now-classic study by Stromswold and colleagues (1996), nearly three decades of neuroimaging studies have looked for syntax in the brain and produced diverse, and at times contradictory, results. A myriad of functional hypotheses has arisen from hemodynamic studies, ranging from hyperlocalisation of syntactic processing in a subpart of the left inferior frontal gyrus (LIFG; Zaccarella & Friederici, 2015) to it relying on an extensive cortical “language network” (Blank et al., 2016; Fedorenko et al., 2020). Electrophysiological techniques like electroencephalography (EEG) and magnetoencephalography (MEG), with their high temporal resolution, have further established that syntax is better construed as a collection of parsing processes than as one uniform operation. In the context of EEG, for example, violation paradigms have identified different syntax-related components based on their latency, duration, and spatial distribution and ascribed them distinct functions, typically beginning with the early left anterior negativity (ELAN) reflecting local phrase structure construction and ending with the late P600 as an index of sentence-level integration processes (see, e.g., Friederici (2002, 2011) for reviews). A complementary strand of research investigating grammatically licit constructions using MEG has corroborated the notion of syntax as a diverse phenomenon. For example, semantically similar expressions of differing syntactic complexity produce early activity increases in the left posterior temporal lobe (LPTL; Flick & Pylkkänen, 2020; Matar et al., 2021) while long-distance dependencies have been found to modulate LIFG activity at considerably later latencies (Leiken & Pylkkänen, 2014; Leiken et al., 2015). As such, the question of *when* and *where* we might find syntax in the brain has required substantial finetuning to adequately capture the spectrum of hypothesized parsing processes.

Despite the methodological variability in how the extant literature has investigated distinct syntactic processes, most studies rely on serially unfolding language, either auditorily in the form of natural speech or visually using the popular stimulus delivery technique Rapid Serial Visual Presentation (RSVP) delivering words one-by-one. Rapid Parallel Visual Presentation (RPVP; Snell & Grainger, 2017), a novel stimulus delivery technique in which stimuli are flashed in their entirety for a few hundred milliseconds, is gaining traction as an alternative approach to study linguistic computations. Unlike serially unfolding language, RPVP imposes no order-processing constraints and can therefore reveal the extent to which our brains engage in parallel perception of language when the stimulus allows it. The key finding from this paradigm is the elicitation of the so-called Sentence Superiority Effect, a phenomenon that was first reported by James Cattell in 1886 and refers to the faster and more accurate recognition of structured vis-à-vis unstructured representations. Through the use of RPVP, a number of studies have elicited behavioural instances of this Sentence Superiority Effect (e.g., Massol et al. (2021); Pegado et al. (2021); Snell & Grainger (2017)), typically by comparing grammatical sentences to their scrambled counterparts. Using EEG, Wen et al. (2019, 2021a) were the first to address the neural signature of the Sentence Superiority Effect as it evolves over time, finding the sentence vs. scrambled words-contrast to manifest as an N400-type effect. Specifically, they observed reduced N400 amplitudes for grammatical sentences compared to scrambled ones from 274-410ms, which they interpreted as a facilitation effect guided by the interactive processing of word identities and a rapidly generated syntactic representation.

Recently, the RPVP studies by Wen and colleagues have been complemented by an additional EEG study (Dunagan et al., 2025) and a few MEG studies (Dufau et al., 2024; Fallon & Pylkkänen, 2024; Flower & Pylkkänen, 2024). Focusing on the MEG studies, they varied in their approach for source localisation. Some employed liberal search areas such as a broadly defined language mask (left temporo-parietal lobes including select frontal areas; Fallon & Pylkkänen, 2024) or the entire left hemisphere (Flower & Pylkkänen, 2024) while Dufau and colleagues (2024) performed targeted regions of interest (ROI) analyses. Each study obtained slightly different but still largely overlapping results. Flower & Pylkkänen (2024) observed two large left fronto-temporal clusters (213–469ms, with a three-way distinction between grammatical sentences, inner-transposed sentences, and reversed sentences from 213–320ms; 448–725ms, with a two-way distinction for grammatical and inner-transposed sentences vs. reversed sentences across the entire cluster) while Fallon & Pylkkänen’s (2024) effects were earlier and included the same middle-posterior temporal regions as well as some parietal regions (sentences vs. noun lists: 127–214ms; 200–259ms). Common to both studies, however, is that they found the neural instances of the Sentence Superiority Effect to manifest as increased activity to structured over unstructured representations. Dufau and colleagues (2024) likewise mostly found increased activity for structured representations, with their ROI analyses yielding clusters in the LIFG (321–306ms; 549–602ms), the left anterior temporal lobe (LATL; 466–531ms), and the left posterior superior temporal gyrus (553–622ms). Despite the different analysis approaches, the effects observed by Dufau and colleagues (2024) converge considerably with those of Flower & Pylkkänen (2024) in terms of time and space. The study by Fallon & Pylkkänen (2024) thus distinguishes itself given the rapidity of the observed effects and the involvement of parietal cortex; however, this can likely be ascribed to the use of noun lists as their non-sentence stimuli instead of scrambled or reversed sentences.

Many of the RPVP studies investigating the Sentence Superiority Effect add additional manipulations, typically varying the type of erroneous stimuli. For example, a number of studies have examined the so-called Transposed-Word Effect, referring to the fact that people do not always detect minor word order-errors as in *you that read wrong* (Flower & Pylkkänen, 2024; Mirault et al., 2018; Pegado et al., 2021; Snell & Grainger, 2019; Wen et al., 2021b). Of particular interest in this case is again the MEG study by Flower & Pylkkänen (2024) in which they, as mentioned above, found sentences with such inner-transpositions to first pattern differently from both grammatical and reversed sentences before patterning similarly to grammatical sentences. To explain this, they proposed that the brain initially detects but subsequently “fixes” the word order-error in the inner-transposed sentences. The Transposed-Word Effect is thus evidence that syntactic structure impacts neural activity almost immediately but that top-down knowledge can override minor errors such as inner-transposed words, suggesting that the brain distinguishes between different levels of ungrammaticality based on the severity of the violation. Other types of erroneous stimuli that have been used to successfully induce instances of the Sentence Superiority Effect include agreement errors (Dunagan et al., 2025; Fallon & Pylkkänen, 2024) and semantic role reversals (Fallon & Pylkkänen, 2024); interestingly, double violation stimuli with both agreement errors and semantic role reversals do not elicit instances of the Sentence Superiority Effect. At present, Fallon & Pylkkänen are the only ones to have investigated a type of grammatical but non-canonical stimuli, namely relative clauses like *wounds nurses clean*, using RPVP. They did not observe an instance of an early neural Sentence Superiority Effect with this type of stimulus. Altogether, the available evidence suggests that at-a-glance processing maps the stimulus to a basic constituent structure in a way that is insensitive to other syntactic properties, such as agreement or displacement.

While the present study also contributes to the characterization of the neural signatures of the Sentence Superiority Effect, notably through the use of minimal two-word stimuli, it goes beyond the relatively coarse contrast of sentences vs. non-sentences and tests the applicability of RPVP to examine different licit linguistic computations at a higher level of granularity. Specifically, we propose using RPVP to reveal aspects of syntactic processing that may be hidden by the serial presentation of language.

### Benefits of parallel presentation as an approach to the neurobiology of syntax

This study takes advantage of two key properties of parallel presentation, which potentially enables a “purer” measurement of syntactic processing than serial presentation. First, when the brain maps a serial input to linguistic knowledge during language comprehension, neural activity is robustly driven by temporally induced incremental predictions of upcoming elements based on previously encountered elements (e.g., Brothers et al. (2015, 2023); Kuperberg et al. (2020); Lau et al. (2006, 2013); Szewczyk & Schriefers (2018); Yano (2018); Zhang et al. (2023)). Since a parallel stimulus does not unfold over time, RPVP allows us to assess linguistic computations in the absence of predictions from a temporally preceding context.

Second, RPVP removes the working memory demands that have been difficult to dissociate from the syntactic processing of dependencies. In research using serial presentation, some studies have deliberately used syntactic dependencies to study syntactic working memory (see, e.g., Christensen et al. (2013); Fiebach et al. (2002, 2005); Makuuchi et al. (2013); Meyer et al. (2013); Santi & Grodzinsky (2007a, 2007b)). More importantly for the present study, however, working memory is a necessary processing component when we understand serially presented syntactic dependencies, and many syntactic dependency manipulations—starting with Stromswold and colleagues’ (1996) seminal study—have also involved differential working memory demands. The issue arises from contrasting sentences like *Whati did the woman win ti?* and *The woman won the marathon*. The former is hypothesised to be syntactically more complex than the latter given the dependency between the *wh*-word and its trace. Specifically, the *wh*-word in *Whati did the woman win ti?* must be held in mind from when encountered in its surface position until integration into its trace position is possible. This necessarily increases the load on working memory—regardless of the exact nature of the content believed to be stored there—compared to a dependency-free sentence like *The woman won the marathon.* As a result, the extent to which heightened activation to structures with syntactic dependencies is not simply due to differential working memory demands has been questioned and, at least by some research, challenged. For example, one line of work has suggested that the commonly observed increase of activation in the LIFG in response to syntactically complex sentences (e.g., Ben-Shachar et al. (2004); Just et al. (1996); Santi & Grodzinsky (2007a, 2007b); Shetreet & Friedmann (2014); Stromswold et al. (1996)) actually reflects articulatory rehearsal (Baddeley et al., 1975), which facilitates the holding in mind of displaced elements (Rogalsky et al., 2008; Rogalsky & Hickok, 2011). According to this hypothesis, the traditional locus of syntax may instead be a phonological working memory hub.

RPVP has the potential to address both challenges. Since a parallel stimulus does not unfold over time, any differential processing of experimental conditions provides a measure of how input is mapped to linguistic knowledge in the absence of predictions from a temporally preceding context. Characterizing the spatiotemporal impact of syntactic dependency manipulations during parallel stimulus presentation will thus be an important step towards a better neurobiological delineation of linguistic computations and prediction. Similarly, eliminating the temporal dimension offers the brain the opportunity to perceive the entire structure at once, which would mitigate the need for storing displaced elements in working memory until they may be integrated into the sentence. Unlike with serial stimulus delivery techniques, the simultaneous presentation of complete sentences thus provides minimal, if any, differential load to working memory across experimental conditions. With its lack of presentation-imposed prediction effects and disparate working memory loads, RPVP might therefore offer a new and exciting window into the study of syntactic processing.

While this study will follow previous RPVP studies in trying to elucidate instances of a Sentence Superiority Effect, the main objective is to use this paradigm to probe for syntactic dependency effects. Specifically, we take advantage of the syntactic properties of Danish to vary the number and type of syntactic dependencies in two-word sentences, each of which are briefly flashed in their entirety during MEG recordings. Our task was a simple matching task in which half of the trials repeated the target stimulus as the task stimulus and half did not, again requiring minimal involvement of working memory (Dunagan et al., 2025; Fallon & Pylkkänen, 2024; Flower & Pylkkänen, 2024). We varied syntactic frame by manipulating the interrogative force of declarative sentences like *pigen festede* (“the girl partied”; Noun-Verb), turning them into yes/no questions like *festede pigen* (“did the girl party”; Verb-Noun) through a mere word order swap. Notably, the surface lexico-semantics therefore stay the same across declarative sentences and yes/no questions unlike in many other studies looking at elements displaced over short and long distances (Gouvea et al., 2010; Leiken et al., 2015; Phillips et al., 2005) as well as when looking at two-word composition with varying degrees of morphological complexity (Matar et al., 2021). As an additional syntactic manipulation, we included verbs that are hypothesised to trigger argument-movement (unaccusative and alternating unaccusative verbs; Perlmutter, 1978) and ones that do not (unergative verbs). The inclusion of two syntactic dependency manipulations allows us to compare any observed effects to each other, with any similarities between them suggesting that they reflect a common underlying operation responsible for resolving syntactic dependencies. To probe for instances of the Sentence Superiority Effect, we contrasted the two-word sentences with two-verb lists as our non-sentence condition. These verb lists paired the intransitive verbs from the sentences with transitive verbs in both word orders (Transitive-Intransitive and Intransitive-Transitive) to emulate the syntactic dependency manipulations of the sentences to the extent possible while still being devoid of hierarchical structure. Notably, two-word sentences are as of yet unexplored in RPVP, making this the first investigation of such minimal structures. Any differential processing of these two-word stimuli—especially if it arises from one of our syntactic dependency manipulations—will provide strong evidence that syntactic structure building is an inherent and deeply automatic trait of the language system, even when encountering rapidly flashed minimal stimuli that on the surface do not require elaborate structures in order to be interpreted.

By using RPVP to study syntactic structure, we chart new territory and—as with any first exploration—there will inevitably be some unknowns. First, it is worth noting that while we have a well-articulated literature of syntactic parsing for serially unfolding language, the theorizing of how the brain reacts when temporal dynamics are not needed is only just beginning. Crucially, this study only allows us to make claims about the syntactic dependency manipulations employed and not, for instance, about the processes involved in parsing more generally. Second, we want to acknowledge that our syntactic dependency manipulations, as with most syntactic phenomena, are intimately intertwined with meaning. As such, the tight control of the surface lexico-semantics in the syntactic frame manipulation is at the expense of different discourse-level interpretations— one being a statement of facts, the other an interrogative—thus making this an inherent confound of this specific manipulation. Similarly, our verb argument structure manipulation may yield effects rooted in semantic rather than syntactic differences given that the different structures of the intransitive verbs also reflect different thematic role assignments. While we will assess the validity of interpreting the effects as syntactic to the extent possible and discuss alternative interpretations when relevant, we maintain an overall syntactic framing of the study. Specifically, we will discuss the results against the backdrop of generative grammar (Chomsky, 1981, i.a.), which assumes generation of all syntactic structure, but they may equally well be accounted for using other theoretical frameworks like tree-adjoining grammar (Joshi et al., 1975; Joshi & Schabes, 1997, i.a.) or construction grammar (Fillmore et al., 1988; Goldberg, 1995, i.a.), both of which posit memorization of smaller chunks that are then combined to larger structures. We chose generative grammar as our framework given the relative sparsity of work on Danish in other theoretical frameworks. The following sections outline the relevant assumptions from generative grammar as well as prior findings from cognitive neuroscience that form the basis of the present investigation.

### Manipulating syntactic frame: An overlooked advantage of V2 languages

Unlike their English counterparts *the girl partied* and *did the girl party*, the corresponding Danish declarative sentence *pigen festede* and yes/no question *festede pigen* contain the exact same lexical material. The ability to manipulate syntactic structure while keeping constant the perceived lexical material was our motivation behind choosing Danish stimuli, and we will therefore briefly sketch which syntactic structures we assume for Danish declarative sentences and yes/no questions and why this is so (Figures 1a-b). Any reference to English below is solely intended to help the reader appreciate the differences between Danish and a language like English, the syntax of which more readers are assumed to be familiar with.

**Figure 1.**
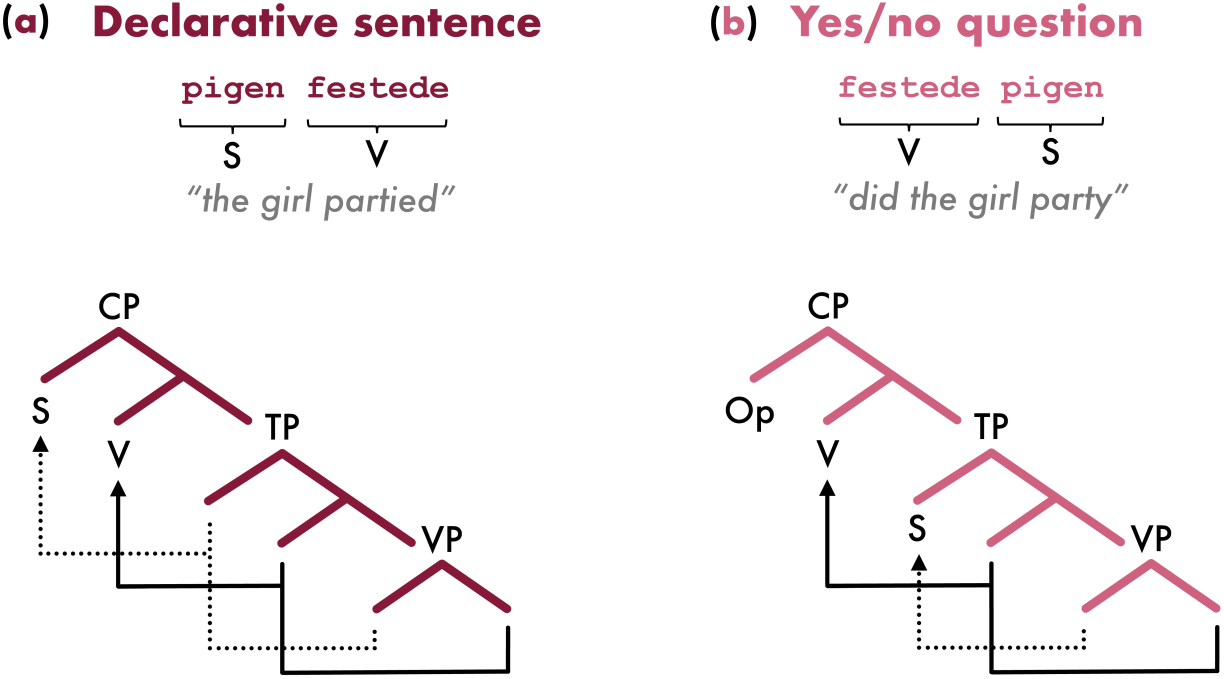
Simplified tree diagrams to illustrate the differences between (**a**) Danish declarative sentences and (**b**) yes/no questions. Arrows indicate displacement of elements from positions lower in the tree (syntactic dependencies). S: subject; V: verb; Op: empty operator; CP: complementiser phrase; TP: tense phrase; VP: verb phrase.

Danish, like most Germanic languages, is a Verb-Second (V2) language. V2 languages are special in that the finite verb is generally the second unit—a so-called constituent—in a clause (originally proposed by den Besten (1983)). While this is also often the case in an SVO language like English, V2 languages differ in that there are few requirements to the first clause constituent, making it possible for various non-subject constituents to occupy this position (see Box 1 for Danish examples). Taking the Danish declarative sentence *pigen festede* as an example (Figure 1a), generative grammar accounts for the flexibility of the first clause constituent by assuming movement of the verb and—in this simple sentence—the subject from their base-generated positions to landing sites high in the syntactic structure (CP). In comparison, the landing sites in non-V2 languages like English are lower. It is precisely the different landing sites of the verbs that allow non-subject constituents to appear before the verb in Danish but not in English. Crucially, the verb and an eligible first-position constituent *must* move to these higher landing sites in V2 languages, even when this additional operation yields the same word order-configuration found prior to movement as is the case in Figure 1a.

Contrary to their declarative counterparts, Danish yes/no questions like *festede pigen* appear to have V1 order rather than V2 order. Such apparent V1 order (also found in imperatives and other exceptional constructions), however, is typically analysed as being covertly V2. This is achieved by merging a covert question operator—a functional element without phonological content— above the verb, thus forcing the subject to stay below it (Christensen, 2008; Holmberg, 2010, 2015; Vikner, 1995; see also Katz & Postal (1964) for non-V2 accounts of covert question operators). Figure 1b illustrates the presumed hierarchical structure for Danish yes/no questions. When comparing Figures 1a-b, it should be clear that although the overall structures of the tree diagrams are comparable, one has more merged elements (yes/no questions) while the other has more syntactic dependencies in the form of displaced elements (declarative sentences).

Despite offering the possibility to compare different syntactic structures while keeping the perceived lexical material constant, experimental studies utilizing such properties of V2 languages remain scarce (but see den Ouden et al. (2008) for a noteworthy exception). The only other neuroimaging study indirectly taking advantage of the syntactic frame manipulation detailed above (Christensen, 2008) does not report results from comparing these particular conditions. If we observe differences in neural activation for the two syntactic frames, this may indicate that building one structure is more costly than building the other. Since merge and move are both assumed to be costly (see, e.g., Grodzinsky et al. (2021) for a recent review) and because we know of no prior studies using a comparable juxtaposition of the two operations, we refrain from making directional predictions about which structure building operation will be the most costly.

### Manipulating verb argument structure: The Unaccusative Hypothesis

Although simple Noun-Verb sentences like *the girl partied* and *the girl awoke*, prima facie, seem structurally similar, the arguments of the verbs have different thematic roles. The argument of *partied* is an agent (the “doer” of the event) and the argument of *awoke* a theme or patient (the entity affected by the event). The original account of why intransitive verbs like *partied* differ from *awoke* was of a syntactic nature, with the different thematic relations being thought to stem from different underlying argument structures (Burzio, 1986; Perlmutter, 1978, i.a.). Whereas the argument of *partied* is a canonical subject, the argument of *awoke* has been hypothesised to originate in the object position before advancing to the subject position. As such, the syntactic derivation of sentences with *awoke*-verbs involves a syntactic dependency not found in sentences with *partied*-verbs. In linguistic theory, this syntactically grounded account is known as the Unaccusative Hypothesis and the two types of intransitive verbs as unergative verbs (*partied*-verbs) and unaccusative verbs (*awoke*-verbs). The structural difference between unergative and unaccusative verbs is believed to explain various linguistic phenomena surrounding resultative constructions (Simpson, 1983), impersonal passivization (Abraham, 1986), and auxiliary selection in a number of Romance and Germanic languages (Haider & Rindler-Schjerve, 1987; Rosen, 1984; Zaenen, 1988). Figure 2 schematises the different argument structures of unergative and unaccusative verbs. So-called alternating unaccusative verbs like *dried*, a subdivision of unaccusative verbs that may optionally be transitive, are also illustrated here.

**Figure 2.**
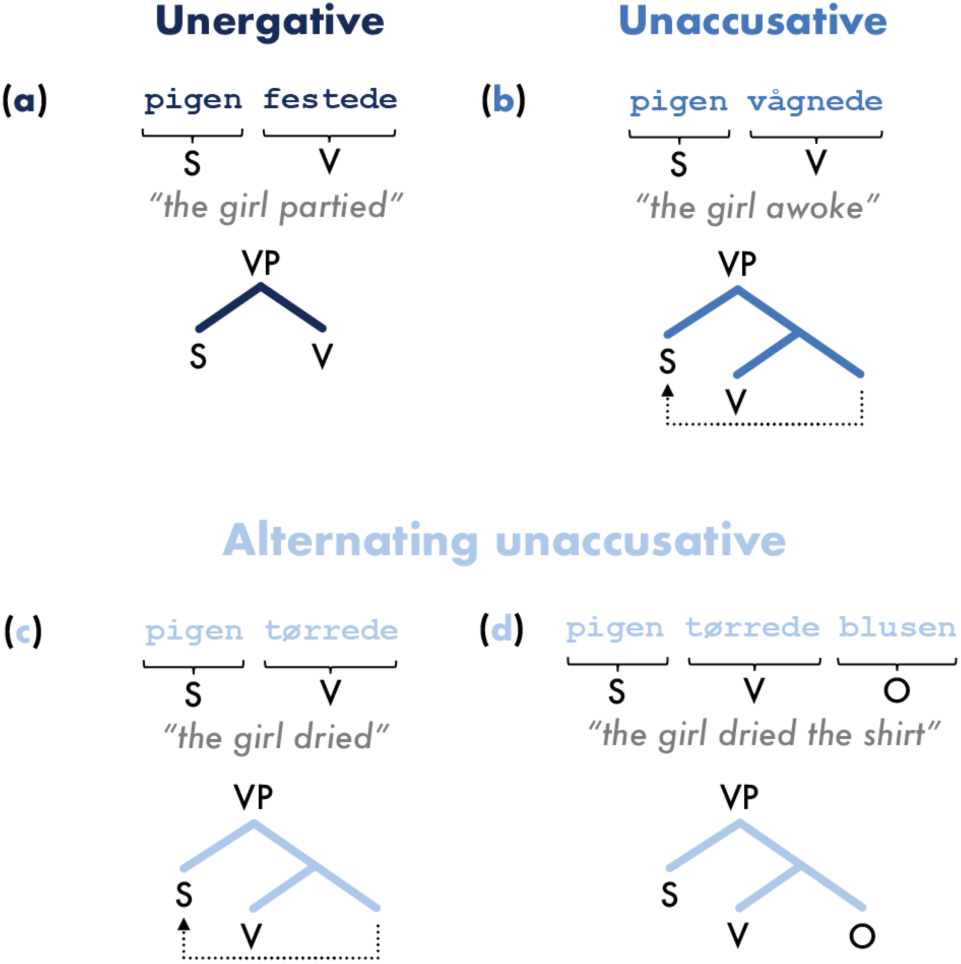
Simplified tree diagrams illustrating the different verb argument structures for (**a**) unergative verbs, (**b**) unaccusative verbs, as well as the (**c**) intransitive and (**d**) transitive forms of alternating unaccusative verbs. Note that all four structures must necessarily be embedded in complete CPs (see Figure 3 for examples). Arrows indicate displacement of elements from positions lower in the tree (syntactic dependencies). S: subject; V: verb; O: object; CP: complementiser phrase; VP: verb phrase.

The distinction between unergative and unaccusative verbs, oftentimes couched within the syntactic framework of the Unaccusative Hypothesis has been extensively researched from a theoretical perspective and has also received experimental support from many methodologies, e.g., behavioural studies with neurotypical and/or aphasic participants (Friedmann et al., 2008; Koring et al., 2012; Koring & Van De Koot, 2018; J. Lee & Thompson, 2011; M. Lee & Thompson, 2004; McAllister et al., 2009; Stavrakaki et al., 2011; but see Huang & Snedeker (2020) for a study finding no difference between unergative and unaccusative verbs), fMRI (Agnew et al., 2014; Meltzer-Asscher et al., 2013, 2015; Shetreet et al., 2009; Shetreet & Friedmann, 2012; but see Wang et al. (2021) for a lack of contrast between unergative and unaccusative verbs), and EEG (Martinez de la Hidalga et al., 2019; Zawiszewski et al., 2022). Generally, unaccusative verbs are found to incur a processing cost compared to unergative verbs as evidenced in higher reaction times, higher error rates, increased neural activation, and a general avoidance of producing constructions with unaccusative verbs. The neural substrates identified as responsible for such asymmetrical processing vary somewhat from study to study. For example, the unaccusative-unergative contrast has been found to implicate as disparate regions as the LIFG (Meltzer-Asscher et al., 2015; Shetreet et al., 2009; Shetreet & Friedmann, 2012), the left superior frontal gyrus (Shetreet et al., 2009), the left middle temporal gyrus (Shetreet et al., 2009; Shetreet & Friedmann, 2012) as well as bilateral superior temporal gyri (Agnew et al., 2014). Similarly, contrasting alternating unaccusative verbs with non-alternating intransitive verbs has sometimes resulted in increased activation in bilateral posterior perisylvian regions as well as bilateral middle and superior frontal gyri (Meltzer-Asscher et al., 2013) and at other times in no differential activation (Meltzer-Asscher et al., 2015).

In this study, we will treat unaccusativity as a syntactic phenomenon by assuming different underlying argument structures for unergative and unaccusative verbs, thus aligning with the Unaccusative Hypothesis. Several syntactic unaccusativity diagnostics are available in Danish and, as described in the *Stimuli* section, we employed stringent selection criteria to only use verbs that can be classified as strictly unergative, unaccusative, or alternating unaccusative. However, different syntactic realizations need not be the whole story of unergative and unaccusative verbs. For example, one influential account by Levin and Hovav (1995) straddles the syntax-semantics interface, acknowledging the syntactic encoding of unaccusativity while emphasizing that unaccusativity is, ultimately, semantically determined. Even more different from the Unaccusative Hypothesis are those accounts explicitly disputing any syntactic grounding and, instead, explaining it solely in terms of semantic properties of the predicate such as agentivity and telicity (Kaufmann, 1995; Shannon, 1992; Van Valin, 1990; Zaenen, 1988, i.a.). Others yet focus less on the exact ontological status of the phenomenon and more on appropriate organization of the different intransitive verbs. Cross-linguistically, some intransitive verbs have eluded binary classification given their variable behaviour across unaccusativity diagnostics, and some researchers therefore propose to abandon binary classification in favour of gradience hierarchies (Baker, 2019; Sorace, 2000).

Common to all the above theories, however, is their entrenchment in linguistic theory; the division of intransitive verbs is contingent on syntactic and semantic constructs that are often hard to operationalise beyond a handful of clear-cut examples. As a recent alternative, Kim et al. (2024) employed an experiential approach, based on Binder et al.s (2016) embodied semantics-model, to chart how the ratings of features belonging to different domains of experience (visual, auditory, motor, social, spatial, temporal, emotion, etc.) correlate with different intransitive verbs. Interestingly, Kim and colleagues (2024) found that their experiential model was better at predicting human naturalness ratings on prenominal past participle-modification, one of the few available unaccusativity diagnostics in English, than models relying solely on syntactic or semantic classification. While the experiential approach has yet to be tested on other combinations of diagnostics and languages, the different experiential content of unergative and unaccusative verbs may go some way in explaining why we observe a seemingly universal cross-linguistic split of intransitive verbs and how children manage to acquire the different types of intransitive verbs, even in languages like English that offer no morphosyntactic clues to unaccusativity.

### The current study

#### Aim 1: The neural indices of syntactic dependencies in a working memory-free paradigm

The main purpose of this study is to investigate how syntactic dependencies—here instantiated through the manipulation of syntactic frame and verb argument structure—are processed neurally in the absence of temporally induced incremental predictions and when condition-specific working memory-related costs are eliminated. If the brain has a unified hub responsible for computing syntactic dependencies, regardless of the type, the same brain region(s) should be responsive to the manipulation of both syntactic frame and of verb argument structure. As such, neural activity may be modulated as a function of the number of movement operations (see Figure 3 for a schematization). Alternatively, the brain may distinguish between different types of syntactic dependencies, resulting in the two manipulations engaging distinct brain regions. A lack of overlap could also mean that the effects stem from non-syntactic operations, which would then require additional investigation. Since this is one of the first investigations of the neural correlates of syntactic dependencies when freed from seriality-induced prediction effects and dissimilar working memory costs across experimental conditions, the effects may be quite different from those identified in prior literature, and we therefore remain agnostic to their localisation in time and space.

**Figure 3.**
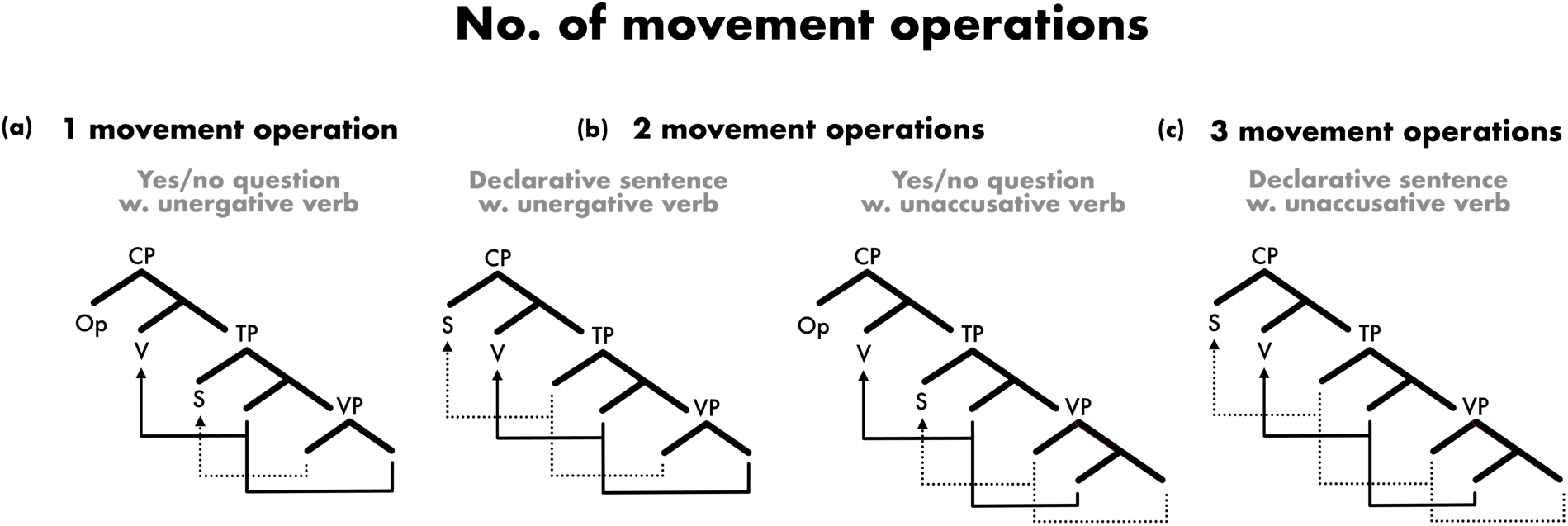
Simplified tree diagrams illustrating the interaction between our syntactic frame and verb argument structure-manipulations, quantified as the number of movement operations between the different conditions. (**a**) Yes/no question with unergative verb (1 movement operation). (**b**) Declarative sentence with unergative verb and yes/no question with unaccusative verb (both 2 movement operations). (**c**) Declarative sentence with unaccusative verb (3 movement operations). Sentences with alternating unaccusative verbs (not depicted) pattern as those with unaccusative verbs. Arrows indicate displacement of elements from positions lower in the tree (syntactic dependencies). S: subject; V: verb; Op: empty operator; CP: complementiser phrase; TP: tense phrase; VP: verb phrase.

#### Aim 2: Parallel presentation of full sentences—the Sentence Superiority Effect in the brain

The use of RPVP as stimulus delivery technique additionally allowed us to investigate—as a secondary research question—the neural bases of the Sentence Superiority Effect that have also been observed in prior RPVP literature. The Sentence Superiority Effect illustrates the impressive speed at which the language system operates when presented with stimuli in their entirety for just a few hundred milliseconds. We anticipate an approximate replication of the timing of the effect reported in prior literature (Dufau et al., 2024; Dunagan et al., 2025; Fallon & Pylkkänen, 2024; Flower & Pylkkänen, 2024; Wen et al., 2019, 2021a), likely with two stages of activation (Flower & Pylkkänen, 2024). Following the dominant trend in the MEG studies using RPVP (Dufau et al., 2024; Fallon & Pylkkänen, 2024; Flower & Pylkkänen, 2024), we further expect it to be realized as increased neural activity to structured over unstructured representations. As for source localization, we again rely on the existing MEG literature and predict that any instances of a neural Sentence Superiority Effect will encompass typical left-lateral language regions, including large portions of the temporal lobe and, potentially, frontal and parietal regions.

## METHODS

### Participants

Twenty-nine Danish-speaking individuals were recruited to participate in the study (14 females, 15 males; median age = 27 years (25^th^ percentile = 24 years; 75^th^ percentile = 32 years) through word-of-mouth and advertising on social media. Since the study was conducted in the New York City area, participants comprised a diverse group including tourists and long-term expats, and information about their language use and proficiency levels was therefore collected using a modified version of the language background questionnaire LEAP-Q (Language Experience and Proficiency Questionnaire; Marian et al., 2007). All participants started acquiring Danish before the age of two and claimed it as their native language, even if some participants had grown up in households where a language other than Danish was also spoken (n = 7). On a 1–10 rating scale, all participants assessed their speaking, listening, and reading skills in Danish to be at ceiling. All participants were neurologically intact with normal or corrected-to-normal vision. Participants provided their informed written consent and were paid $30 for their participation. The study was approved by the Institutional Review Board (IRB) ethics committee of New York University (approval number 2016-91).

### Stimuli

The study employs a 2 × 2 × 3 factorial design varying composition (COMP), syntactic frame (SYN), and verb argument structure (ARG). First, a battery of diagnostics was employed to identify Danish verbs that could be classified as unergative, unaccusative, or alternating unaccusative. The diagnostics were: (i) auxiliary selection in the perfect, (ii) ability to occur with a reflexive pronoun in resultative constructions, (iii) ability to occur in impersonal passive constructions, (iv) ability to form nominal modifiers from the past participle, (v) ability to occur with direct objects, and (vi) ability to undergo passivization. Diagnostics (i-iv) separated unergative verbs from unaccusative and alternating unaccusative verbs (see Levin & Hovav (1995) for a review of these diagnostics) whereas (v-vi) were tests of transitivity, primarily used to distinguish unaccusative verbs (intransitive) from alternating unaccusative verbs (optionally transitive). Since different diagnostics may group verbs into slightly different categories, we chose a conservative approach requiring verbs to align with the requirements of each diagnostic for a given verb category. As such, our goal was to select only verbs that—based on our battery of diagnostics—could be classified as categorically unergative or categorically unaccusative. Verb classification was based on the judgments of the first author, who is a native speaker of Danish, and cross-checked with dictionary definitions (Den Danske Ordbog; DSL, 2021) and corpus frequencies from a Danish corpus (KorpusDK; Asmussen, 2012, 2015). For example, (i) the auxiliary selection-diagnostic refers to the fact that Danish has a split auxiliary system, i.e., some verbs take the auxiliary *være* (“to be”) and others the auxiliary *have* (“to have”) in the perfect (Bentley & Eythórsson, 2004; Bjerre & Bjerre, 2007; Larsson, 2014). The choice of auxiliary is tightly correlated with the unaccusativity of the verb, with unaccusative verbs employing the auxiliary *være*, unergative and transitive verbs employing the auxiliary *have,* and alternating unaccusative verbs employing either auxiliary depending on their transitivity in a given sentence. Here, we determined the appropriate auxiliary for each verb by first consulting the dictionary definitions, which specify whether a verb must obligatorily take either *være* or *have* or if both auxiliaries are available. This information was then ratified using corpus searches to compare the ratios of *være* and *have* used with each verb; unergative and unaccusative verbs were only included if they occurred (almost) exclusively with the appropriate auxiliary whereas alternating unaccusatives were only included if they regularly occurred with both auxiliaries. In cases of doubt, for example if the corpus searches did not yield many occurrences, decisions were made by the first author. Similar processes were repeated for the other diagnostics.

A total of twenty-seven intransitive verbs (7–10 letters long) were chosen as critical verbs, resulting in nine verbs for each of the three categories of ARG. Efforts were made to reflect a wide range of meanings for each category of verbs and to keep lexical characteristics such as number of letters, syllables, morphemes, and phonemes as well as word frequencies relatively similar, although especially the latter proved challenging. Since the small number of verbs in each category makes it difficult to establish normality and therefore precludes reliable statistical testing of differences between the three verb categories, we merely report descriptive statistics for each category of ARG (Table 1). As can be gleaned from the table, unaccusative verbs tended to be slightly longer than both unergative and alternating unaccusative verbs. While full control of lexical characteristics for the three verb types had been desirable, strict control with the argument structure of the verbs was deemed more important for the purpose of the present study.

**Table 1.**
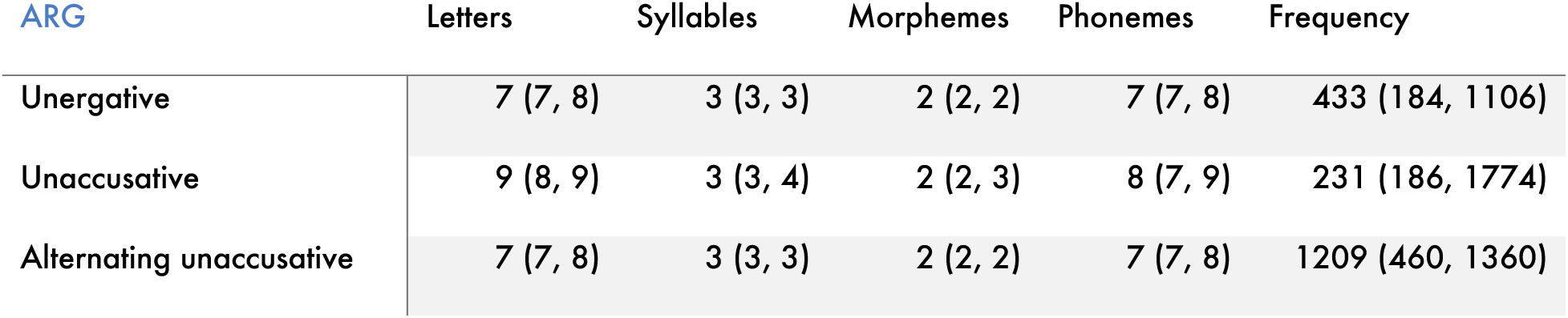
Lexical characteristics of the different verb types. The median value for each measure is reported with the 25^th^ and 75^th^ percentiles in parentheses. ARG = verb argument structure.

| ARG | Letters | Syllables | Morphemes | Phonemes | Frequency |
| --- | --- | --- | --- | --- | --- |
| Unergative | 7 (7, 8) | 3 (3, 3) | 2 (2, 2) | 7 (7, 8) | 433 (184, 1106) |
| Unaccusative | 9 (8, 9) | 3 (3, 4) | 2 (2, 3) | 8 (7, 9) | 231 (186, 1774) |
| Alternating unaccusative | 7 (7, 8) | 3 (3, 3) | 2 (2, 2) | 7 (7, 8) | 1209 (460, 1360) |

Next, each of the twenty-seven intransitive verbs was paired with nine definite nouns. The nouns were 5–7 letters long with generic person reference, e.g., the Danish equivalents of *the girl* or *the man.* All intransitive verbs and nouns were paired in two ways corresponding to the two levels of SYN: Noun-Verb (declarative sentences) and Verb-Noun (yes/no questions). Crucially, the Verb-Noun word order unequivocally produces a yes/no question and, in line with standard practices regarding the use of punctuation in experimental research, we chose to not include a question mark for the yes/no questions. All verbs were in the past tense due to the homonymous nature between many Danish nouns and present tense verbs, and past tense verbs thus enforced compositional rather than list interpretations of the sentences. The transition probability between the two word orders for the nouns and verbs in question, as calculated using KorpusDK, did not differ significantly (*p* = 0.2108). While this metric is somewhat conflated since the Verb-Noun order is also found with preposed non-subject constituents, it suffices to dispel interpretations hinging on word category-based structural frequency effects. In summary, 486 sentences were constructed by pairing each of the twenty-seven intransitive verbs with each of the nine nouns in the two different word orders, meaning that each verb was repeated eighteen times across the sentence stimuli.

Finally, a control condition with two-word lists was created by replacing the nouns in each of the 486 sentences with nine transitive past tense verbs. The transitive verbs matched the nouns in length (5–7 letters) and passed all diagnostics outlined in Box 2. Verb lists were preferable to noun lists as they effectively mitigated any unintended composition effects arising from two-word noun lists (e.g., due to noun-noun compounding or the aforementioned homonymous nature between many Danish nouns and present tense verbs). The complete stimulus set thus had 972 trials. Each stimulus subtended a horizontal visual angle between 4.8° and 6.6° (mean ± SD visual angle: 5.4° ± 0.5°; calculated using an average eye-to-screen distance of 44cm). See Figure 4a for the complete factorial design.

**Figure 4.**
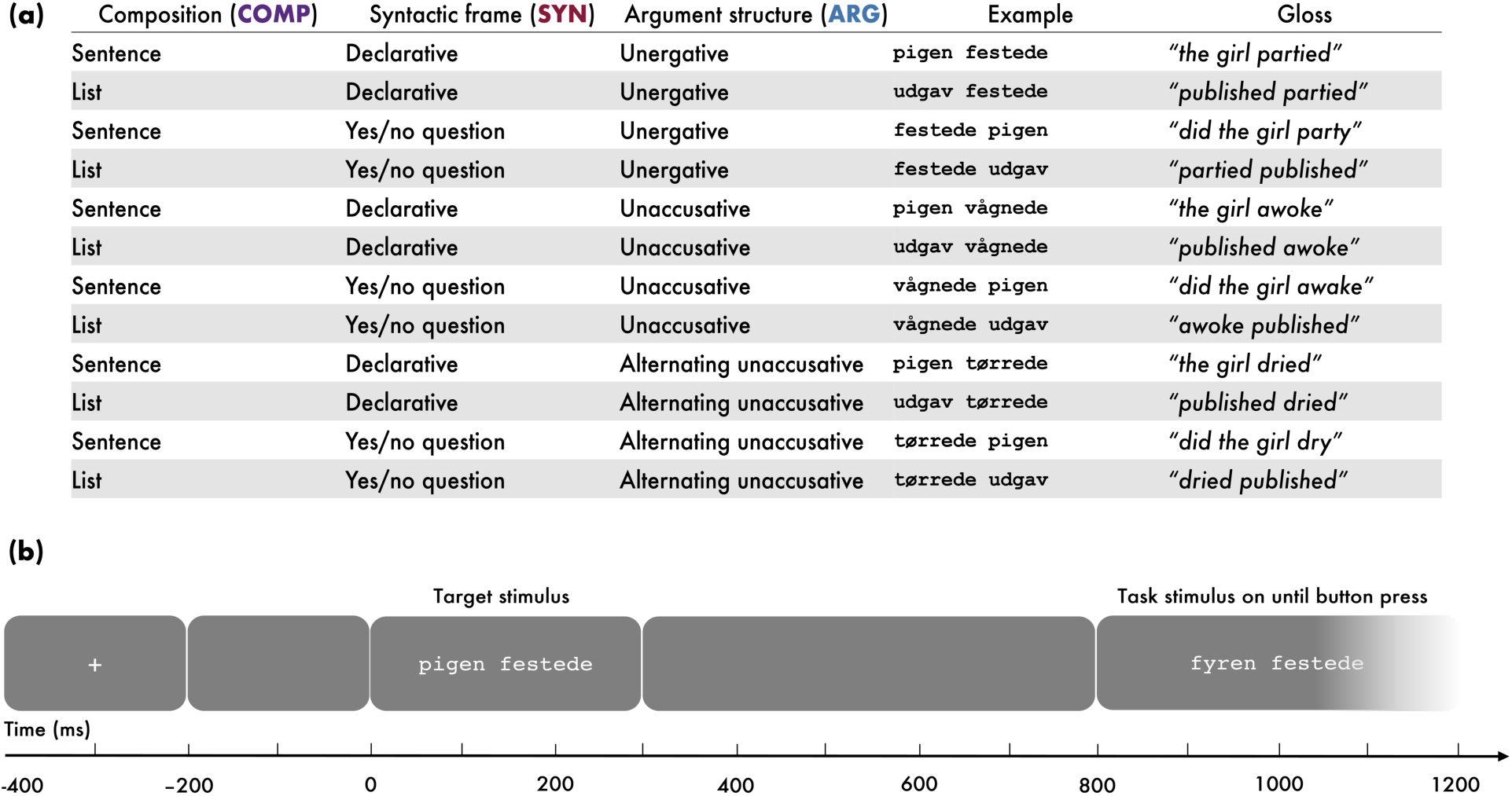
Design table and trial structure. (**a**) Full factorial design with factors composition (COMP), syntactic frame (SYN), and verb argument structure (ARG) alongside example trials for each condition. (**b**) Trial structure illustrating a mismatch trial; the target stimulus and task stimulus differ from each other by one word. The width of the grey boxes indicates length of time on screen.

### Task rationale and trial structure

While the majority of studies using RPVP have employed grammaticality judgments or post-cued partial report, we opted for a straightforward matching task in which participants identified whether target and task stimuli were the same (a match trial) or different (a mismatch trial) (see also Dunagan et al., (2025), Fallon & Pylkkänen (2024); Flower & Pylkkänen (2024), and Pegado et al. (2021) for similar tasks). Main motivations for the task were its speed and the absence of any explicit judgments on the stimuli. Since the task can be performed purely perceptually, any effects of our structural manipulations would be reflective of truly automatic processing. In our study, only one of the two words differed between the target and task stimuli, with swapped words always being of the same length and belonging to the same category (noun, transitive verb, intransitive verb). The identity and position of the swapped words were randomised across the 972 trials. Note that only responses to the first sentence of a trial (the target stimulus) and not the second sentence (the task stimulus) were analysed.

The structure of each trial was as follows (see Figure 4b). First, a fixation cross was displayed (200ms) followed by a blank screen (200ms). Next, the target stimulus was briefly flashed (300ms) and then followed by another blank screen (500ms). Lastly, the task stimulus was displayed until the participant, via button-press, indicated the trial as being either a match or a mismatch trial. Behavioural reaction times were measured from the presentation of the task stimulus. The intertrial stimulus interval was 600–750ms, jittered in increments of 50ms. The 972 trials were randomly divided into nine blocks of 108 trials each and presented centrally on a screen in Courier New-font using PsychoPy 2022.2.4 (Peirce et al., 2019).

### Procedure

The experiment took place at the NYU/KIT MEG LAB at New York University, New York. Upon arrival to the lab, participants gave informed consent and filled out modified versions of a handedness questionnaire (Oldfield, 1971) and the LEAP-Q on paper. Next, participants’ head shapes were 3D digitised using a Polhemus FastSCAN laser scanner (Polhemus, VT, USA) to enable coregistration of the MEG data with the FreeSurfer fsaverage brain (Fischl, 2012) during analysis. Five future marker coil placements and the position of three fiducial landmarks (nasion, left tragus, right tragus) were included in the digitizations. The head shape-scans were followed by a brief instruction period and a practice session of 12 trials mirroring the actual experiment in structure and task but with different stimuli. Participants were then guided into the dimly lit magnetically shielded room hosting the MEG machine where marker coils were placed on the digitised points before the participants completed the experiment in supine position. Each experimental session took ∼2 hours.

### MEG data acquisition and preprocessing

Continuous MEG data were recorded using a whole-head, 157-channel axial gradiometer system (Kanazawa Institute of Technology, Kanazawa, Japan) at a sampling rate of 1000Hz with online high- and low-pass analogue filters of respectively 1Hz and 200Hz. Participants’ head positions relative to the MEG sensors were recorded using marker coils before and after the experiment.

The MEG data were first noise-reduced based on data collected from reference channels using the Continuously Adjusted Least-Squares Method (Adachi et al., 2001) in the MEG160 software (Yokogawa Electrical Corporation and Eagle Technology Corporation, Japan). All other preprocessing steps were conducted using the MNE-Python v0.23.4 (Gramfort et al., 2014) and Eelbrain v0.37.6 (Brodbeck et al., 2022) packages in the Python computing environment. First, an offline 1–40 Hz digital bandpass filter was applied, and flatlined or clearly malfunctioning channels (averaging 3.8 channels per participant) were removed. Epochs were then extracted from 100ms before to 800ms after the onset of the target stimulus; as such, only processing of the target stimulus—not the task stimulus—was included in an epoch. Afterwards, Independent Component Analysis (ICA) was performed to remove artifacts such as eye blinks, saccades, heartbeats, and well-characterised external noise sources. Individual epochs were rejected if they contained amplitudes exceeding 3000 fT or, upon visual inspection, contained sudden increases in the magnitude of the signal (as caused by, e.g., muscular movements), resulting in the removal of 1.91% of epochs. Finally, epochs with incorrect responses were removed and the number of epochs balanced across conditions, leaving on average ∼90.29% of epochs for each participant after epoch rejection and equalization. All epochs were baseline corrected using the 100ms before the onset of the target stimulus.

Dynamical Statistical Parameter Mapping (dSPM; Dale et al., 2000) was used to create cortically constrained estimates of source-level activity for the analyses. During coregistration, the digitised head shape and fiducial landmarks of each participant were scaled and mapped to the Freesurfer fsaverage brain in the creation of cortical surfaces. From these, two source spaces of 2,562 sources (one per hemisphere) were generated and, using the Boundary Element Model (BEM) method, a forward solution computed. A noise covariance matrix was then calculated using the first 100ms of the epochs, which, in tandem with the forward solution, formed the basis for computing inverse solutions with free orientation. Lastly, the noise-normalised minimum norm estimates were converted to dSPMs, enabling spatiotemporal visualizations of activity.

### Statistical analyses

#### Behavioural data

In order to test for behavioural correlates of the Sentence Superiority Effect and our syntactic manipulations, we first cleaned the data of responses more than 3 SDs from overall participant or item means. Descriptive statistics reported in the *Results* section, e.g., means and standard deviations, are calculated on this cleaned data. Then we fitted a total of four models to the data. In all four models, we generated *p* values via likelihood ratio tests and corrected pairwise comparisons using a Tukey adjustment. The analyses were conducted using the *lme4* (Bates et al., 2015) and *afex* (Singmann et al., 2023) packages in R (v4.3.1) and RStudio (v2023.06).

Our first two models probed for instances of the Sentence Superiority Effect, which we should see behavioural correlates of given prior literature, using the full factorial design. Specifically, we fitted a linear mixed-effects regression model to log-transformed reaction time data for the correct responses with COMP, SYN, ARG, and task type (TASK; match/mismatch trial) as fixed effects alongside all possible interaction terms. TASK was included as a fixed effect in all behavioural analyses due to the expectation that match and mismatch trials would asymmetrically recruit cognitive resources; match trials involved priming between target and task stimulus while mismatch trials did not. The maximal random-effects structure that converged included by-participant and by-item varying intercepts. Accuracy data were analysed using a generalised linear mixed-effects logistic regression model (incorrect responses included) with the same structure of fixed effects but with just by-participant intercepts. In both models, we treat significant main effects of COMP as evidence of Sentence Superiority. However, we are also interested in interaction effects involving COMP and significant pairwise comparisons for SYN-ARG combinations since such condition-specific instances of the Sentence Superiority Effect may contribute to our understanding of what drives the Sentence Superiority Effect.

Then, in a more exploratory vein since our focus is on neural rather than behavioural correlates of our syntactic manipulations, we dropped all list stimuli and fitted two models (one for reaction time data, one for accuracy data) to the sentence stimuli to test for effects of our syntactic manipulations. While this could also have been done using the full factorial design—in so doing looking for effects of SYN or ARG modulated by COMP—we chose to separate sentences from lists based on the undertheorised nature of the computations underlying two-verb list processing. These two reduced models were identical to the models using the full dataset except that COMP was no longer included as a fixed effect. As such, these exploratory behavioural analyses focus on effects of SYN and ARG, regardless of whether they manifest as main or interaction effects.

#### MEG data

The analyses of the MEG data were largely exploratory given the use of RPVP, a stimulus delivery technique novel to MEG research at the time of data analysis, which allowed us to investigate the neural indices of the Sentence Superiority Effect and of syntactic dependencies freed from differential working memory costs across experimental conditions. As such, we performed nonparametric spatiotemporal cluster tests (Maris & Oostenveld, 2007) in liberal search areas to maximally capture the effects of our manipulations.

First, we fitted a 2 × 2 × 3 repeated measures ANOVA with factors COMP (sentence, list), SYN (declarative sentence, yes/no question), and ARG (unergative verb, unaccusative verb, alternating unaccusative verb) to the data in bilateral full-hemisphere searches across the entire epoch (0–800ms) to assess instantiations of a Sentence Superiority Effect. Next, to probe for effects of our syntactic manipulations, we dropped all list stimuli and performed 2 × 3 repeated measures ANOVAs with factors SYN and ARG on the sentence stimuli only, thus mirroring the behavioural analyses. Our primary analysis path was to extract the spatial distribution of any clusters associated with instances of a Sentence Superiority Effect as functional regions of interest (fROIs) and perform searches within these in early (100–500ms) and late (500–800ms) time windows. However, since we did not know if any such would arise, we also planned to conduct bilateral lobe-by-lobe searches in these same time windows. The more subtle contrasts in the levels of SYN and ARG compared to COMP motivated the use of these narrower spatiotemporal search areas. In terms of temporal delineation, we opted for time windows that could capture common EEG components like the N400 and the P600 respectively. As to spatial delineation, we chose the more generous search areas of lobes over narrower regions identified in prior literature on syntactic processing, e.g., the LIFG or the LPTL, since parallel stimulus presentation might engage brain regions different from those recruited during serial stimulus presentation. As a reviewer pointed out, the lobe-by-lobe stratification is not a conventional approach but was chosen as a compromise solution, allowing us to stay open for what results might look like given the novelty of RPVP while simultaneously making it possible for us to capture effects of our relatively subtle syntactic manipulations. For completeness, we also performed full-hemisphere analyses across the entire epoch.

All spatiotemporal permutation clustering tests were performed in Eelbrain v0.39.10 (Brodbeck et al., 2023) using a standard procedure. An uncorrected repeated-measures ANOVA was fitted to each spatiotemporal dimension ((f)ROI/time window) to calculate *F* values for source-time point combinations. Clusters were computed from adjacent spatial and temporal values when these met the cluster-forming threshold of *p* < 0.05. Then, cluster *F* values were summed if the cluster consisted of a minimum of 20 vertices and 20 contiguous time points, yielding a cluster-level statistic. We determined significance using a cluster-based (Monte Carlo) permutation test, generating a null distribution of the test statistic for each spatiotemporal dimension by randomly shuffling condition labels for each participant 10,000 times. The observed clusters were then compared to the null distribution and assigned corrected *p* values that reflected the proportion of permuted clusters with sizes larger than the actually observed cluster. In the lobe-by-lobe analyses, the resulting *p* values were corrected for multiple comparisons across ipsilateral lobes (i.e., a correction factor of 4) by controlling the false discovery rate at the critical value of 0.05 (Benjamini & Hochberg, 1995). Given the uncorrected nature of the clusters initially chosen for analyses, the borders of the reported clusters should be interpreted as approximate, thus making it impossible to make exact claims about latency or duration of any effects (Sassenhagen & Draschkow, 2019).

## RESULTS

### Behavioural results

#### Behavioural instances of the Sentence Superiority Effect for reaction times and accuracies

As shown in Figures 5 and 6c, we found clear behavioural instances of the Sentence Superiority Effect, our primary focus of the behavioural analysis. This manifested as significant main effects of COMP (*p* < 0.001) across both reaction time and accuracy data. When following up on these main effects of COMP with post-hoc pairwise comparisons, almost all SYN-ARG combinations yielded condition-specific instances of the Sentence Superiority Effect for reaction time data across all trials, match trials, and mismatch trials while this was only the case for certain SYN-ARG combinations in the accuracy data. Such asymmetry between reaction times and accuracy is expected given the short stimuli and the simple nature of the task, which also resulted in very high overall task accuracies (mean ± SD: 96.42 ± 2.61%) and low reaction times on correct trials (679.6 ± 228.6ms). The following will focus on the effects of COMP and their interactions; however, all likelihood ratio test results for the two models using the full factorial design on both sentence and list stimuli are available in Table 2. Figure 5 illustrates—across the entire factorial design—the significant effect of COMP on reaction times for (a) all trials, (b) match trials, and (c) mismatch trials and on accuracies for (d) all trials, (e) match trials, and (f) mismatch trials. Main effects of COMP averaged over SYN, ARG, and TASK are depicted in Figure 6c.

**Figure 5.**
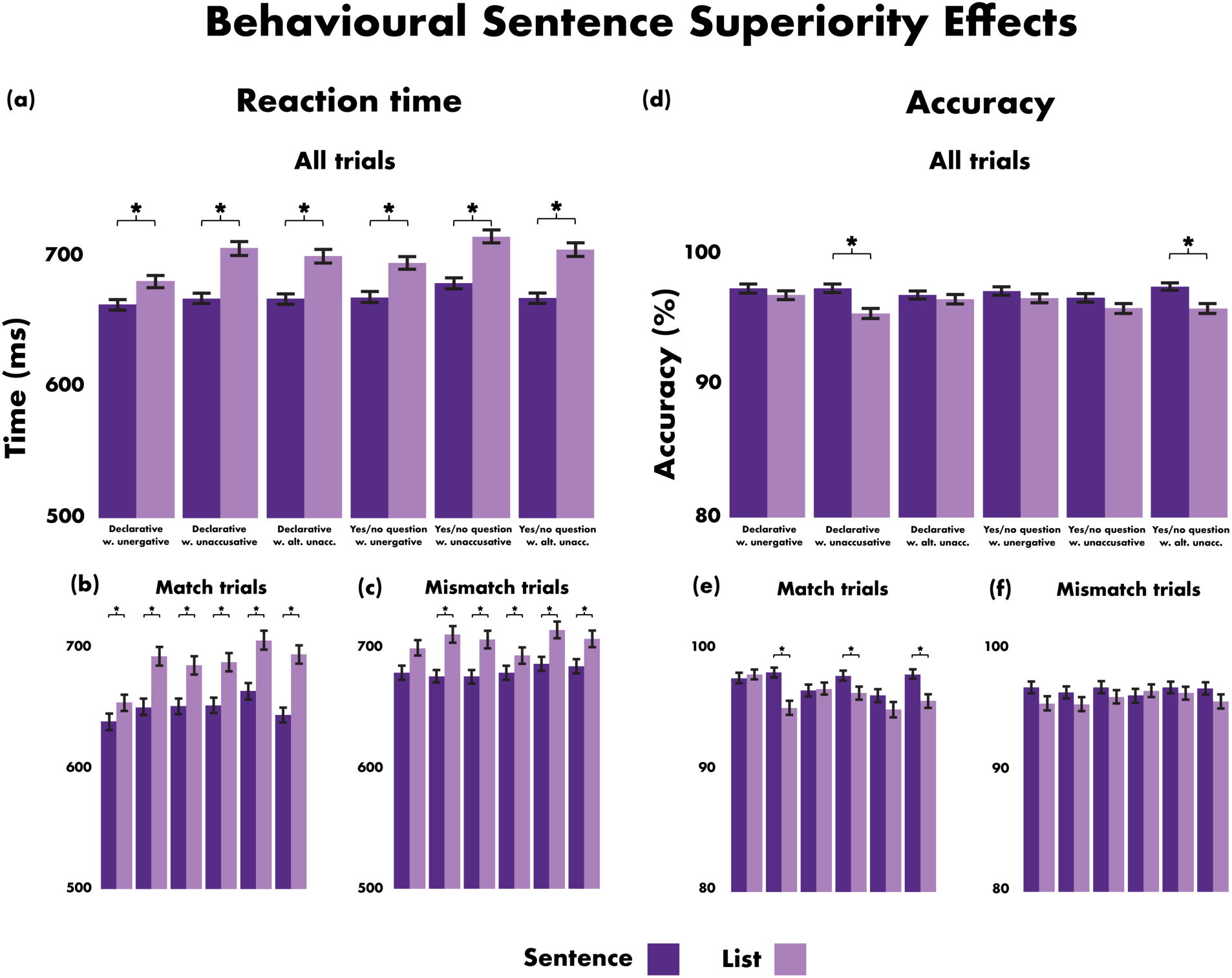
Instances of a behavioural Sentence Superiority Effect plotted for the entire factorial design. On the left, an instance of the behavioural Sentence Superiority Effect for reaction times across (**a**) all trials, (**b**) match trials, and (**c**) mismatch trials. On the right, an instance of the behavioural Sentence Superiority Effect for accuracies across (**d**) all trials, (**e**) match trials, and (**f**) mismatch trials. While COMP was involved in a main effect as well as significant three- and four-way interaction effects for the accuracy data, driven partly by the significant pairwise comparisons indicated by asterisks in the plot, COMP only arose as a main effect for the reaction time data. As such, the statistical significance of pairwise comparisons for SYN-ARG combinations in the reaction time data indicate our post-hoc characterization of the main effect. COMP: composition; SYN: syntactic frame; ARG: verb argument structure.

**Figure 6.**
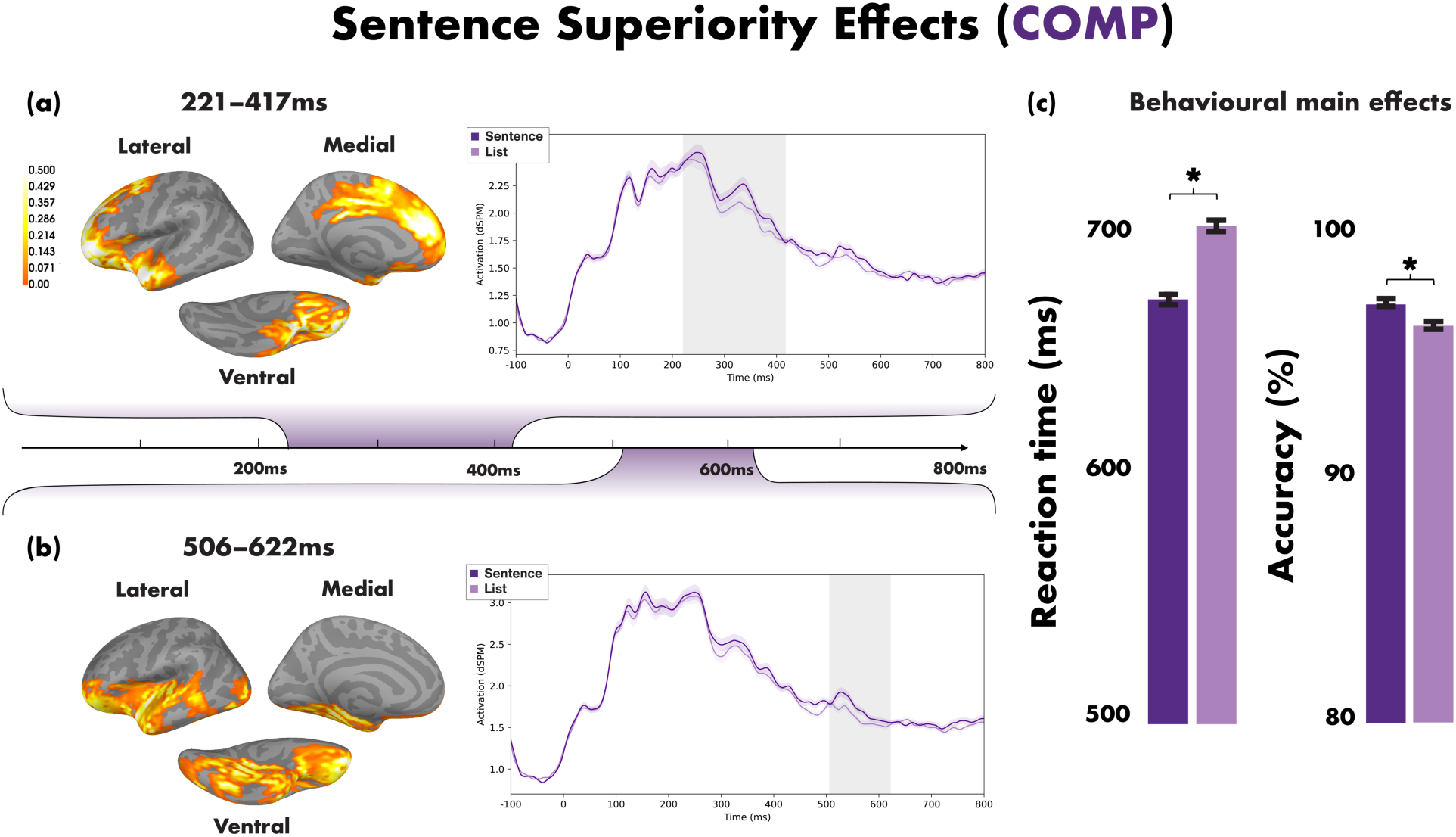
Timeline showing neural effects of manipulating COMP, with the time course for each effect plotting the averaged activity in the identified spatial cluster and the light grey-shading indicating the extent of the significant clusters. (**a)** The first instance of the Sentence Superiority Effect. Neurally, it manifested with a broad fronto-temporal distribution in the left hemisphere from ∼220–415ms post stimulus onset, with increased activity to sentences over lists. (**b)** The second instance of the Sentence Superiority Effect from ∼50–620ms. This cluster likewise showed increased activity to sentences over lists but also included more posterior-ventral temporal areas. (**c**) Behavioural instances of the Sentence Superiority Effect for reaction time and accuracies. COMP: composition; * = p < 0.05 corr.

**Table 2.** Results of likelihood ratio tests for linear mixed-effects regression models for reaction times and accuracies using the entire dataset (sentences and lists). COMP: composition; SYN: syntactic frame; ARG: verb argument structure; TASK: match/mismatch trial.

|  | df | Reaction time |  | Accuracy |  |
| --- | --- | --- | --- | --- | --- |
| | | $\chi^2$ | p value | $\chi^2$ | p value |
| COMP | 1 | 108.78 | < 0.001 | 17.73 | < 0.001 |
| SYN | 1 | 7.98 | 0.005 | 0.58 | 0.447 |
| ARG | 2 | 20.81 | < 0.001 | 5.20 | 0.074 |
| TASK | 1 | 290.84 | < 0.001 | 7.52 | 0.006 |
| COMP × SYN | 1 | 0.08 | 0.784 | 0.09 | 0.766 |
| COMP × ARG | 2 | 5.64 | 0.060 | 2.21 | 0.331 |
| COMP × TASK | 1 | 4.68 | 0.030 | 1.69 | 0.193 |
| SYN × ARG | 2 | 3.22 | 0.200 | 0.69 | 0.707 |
| SYN × TASK | 1 | 10.63 | 0.001 | 2.93 | 0.087 |
| ARG × TASK | 2 | 3.71 | 0.156 | 4.32 | 0.115 |
| COMP × SYN × ARG | 2 | 0.68 | 0.713 | 6.11 | 0.047 |
| COMP × SYN × TASK | 1 | 1.78 | 0.183 | 1.96 | 0.162 |
| COMP × ARG × TASK | 2 | 0.19 | 0.911 | 1.61 | 0.448 |
| SYN × ARG × TASK | 2 | 5.10 | 0.078 | 4.26 | 0.119 |
| COMP × SYN × ARG × TASK | 2 | 0.28 | 0.867 | 6.72 | 0.035 |

Overall, the significant effects of COMP and TASK patterned as expected. Responses were faster and more accurate on sentences (664.3 ± 208.4ms; 96.89 ± 2.33%) than lists (695 ± 246.5ms; 95.95 ± 2.99%), providing instances of the Sentence Superiority Effect. Similarly, responses to match trials were faster and more accurate (667 ± 238.3ms; 96.65 ± 3.12%) than to mismatch trials (692.1 ± 217.8ms; 96.16 ± 2.80%). Furthermore, as suggested by the significant interaction effect between COMP and TASK for reaction times, the extent to which one effect modulated the other varied. As such, although pairwise comparisons revealed significant Sentence Superiority Effects for both match and mismatch trials in isolation (both *p* < 0.001), it was more pronounced for match trials (average difference in reaction time between sentences and lists: 36.4ms) than mismatch trials (25.2ms). Similarly, while sentences and lists each had significantly faster reaction times for match trials compared to mismatch trials (both *p* < 0.001), pairwise comparisons found the impact of TASK to be greater for sentences (average difference in reaction time between match and mismatch trials: 30.8ms) than for lists (19.6ms).

For the accuracy data, COMP was also involved in a three-way interaction effect between COMP, SYN, and ARG and a complete four-way interaction effect. Pairwise comparisons revealed these to be driven partly by instances of the Sentence Superiority Effect for specific SYN-ARG combinations (three-way interaction effect: declarative sentences with unaccusative verbs (*p* = 0.003) and yes/no questions with alternating unaccusative verbs (*p* = 0.0023); four-way interaction effect: declarative sentences with unaccusative verbs in match trials (*p* = 0.002), yes/no questions with unergative verbs in match trials (*p* = 0.0459), and yes/no questions with alternating unaccusative verbs in match trials (*p* = 0.0053)). Given the overall high accuracies and the lack of any clear hypothesis that can account for this pattern, we refrain from further discussion of why only these particular SYN-ARG combinations participate in the interaction. For the reaction time data, the only significant interaction effect involving COMP was with TASK as described above.

#### A syntactic frame effect for sentences with unaccusative verbs in match trials

Our exploratory analysis revealed few interesting effects of our syntactic dependency manipulations in the behavioural data; however, this is not surprising given the subtle contrasts in the levels of SYN and ARG compared to COMP. Table 3 reports all likelihood ratio test results for the reduced models using only the sentence stimuli. Other than reiterating the significant effects of TASK, the only other significant effect was an interaction between SYN, ARG, and TASK in the accuracy data. Pairwise comparisons revealed this effect to be driven partly by an effect of SYN in the case of sentences with unaccusative verbs in match trials (declarative sentences were significantly more accurate than yes/no questions: *p* = 0.008).

**Table 3.**
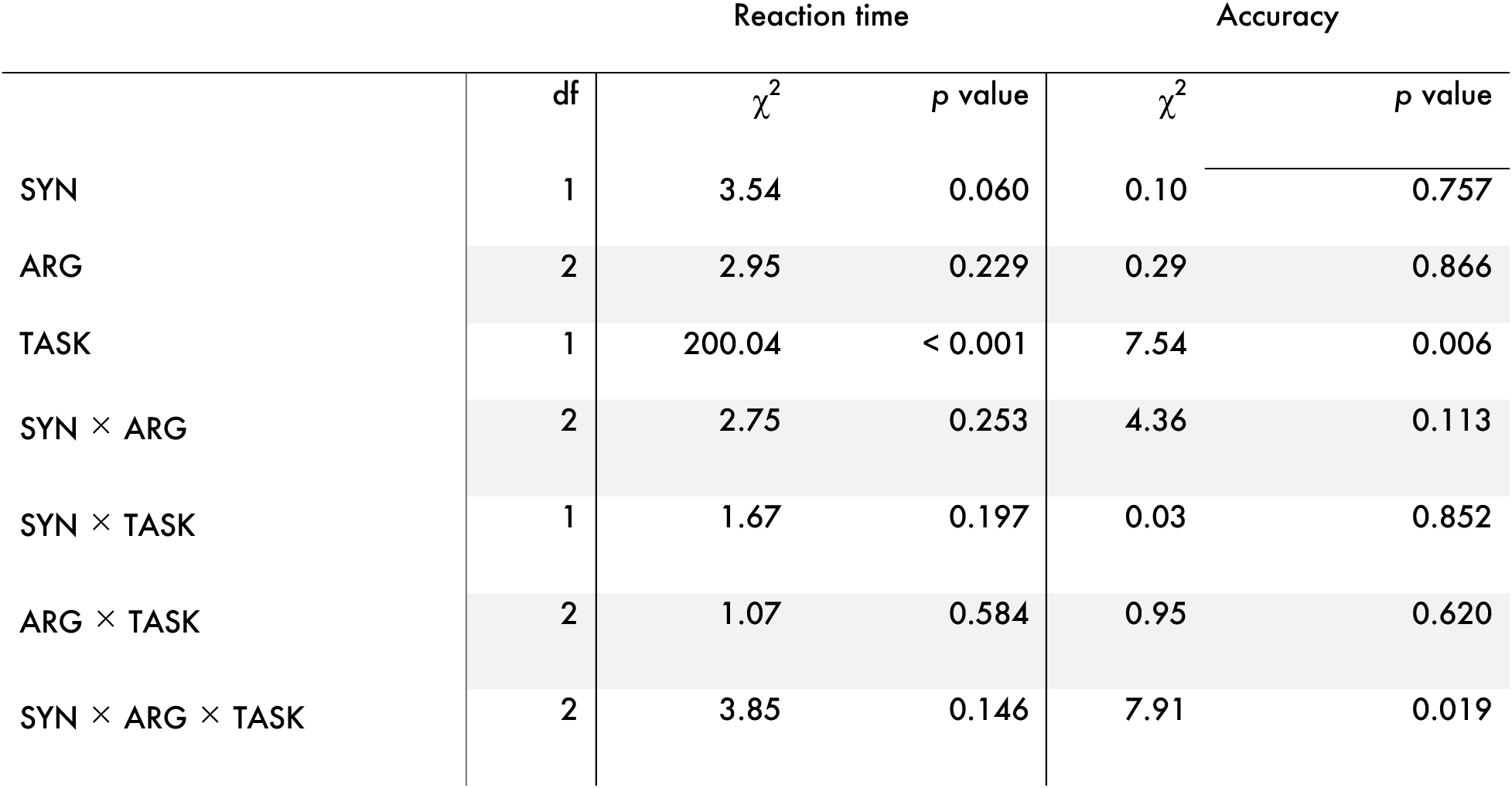
Results of likelihood ratio tests for linear mixed-effects regression models for reaction times and accuracies using only the sentence stimuli. SYN: syntactic frame; ARG: verb argument structure; TASK: match/mismatch trial.

While the undertheorised nature of two-verb lists is our rationale for discarding the list stimuli in the analyses of our syntactic dependency manipulations, we did observe a significant main effect of ARG in the full model of reaction time data (*p* < 0.001) of which there was no trace in the reduced model of reaction time data (*p* = 0.229). Post-hoc decomposition of the ARG effect from the full model reveals it to be driven by list stimuli (faster reaction times to unergative verbs compared to unaccusative (*p* < 0.001) and alternating unaccusative verbs (*p* = 0.018); no significant difference in reaction times to unaccusative and alternating unaccusative verbs (*p* = 0.079)) rather than sentence stimuli (*p* > 0.24 for all pairwise comparisons). Although our manipulation of ARG was conceptualised as varying the number of syntactic dependencies in a sentence, the three verbs may be partitioned in the same way based on their lexico-semantic characteristics. In other words, it is as likely that sentences with unergative verbs are different from sentences with unaccusative or alternating unaccusative verbs because they differ in the number of syntactic dependencies as it is due to different thematic role assignment. Notably, however, such conflation of syntax and semantics is not found for the list stimuli, which are presumed to be devoid of syntactic structure. The difference in reaction times between verb types when embedded in lists is thus likely to be of a lexico-semantic, not syntactic, nature.

### MEG results

#### Early and late fronto-temporal instances of a Sentence Superiority Effect in the left hemisphere

To uncover any neural correlates of the Sentence Superiority Effect, we ran a 2 (COMP: sentence, list) × 2 (SYN: declarative sentence, yes/no question) × 3 (ARG: unergative verb, unaccusative verb, alternating unaccusative verb) repeated-measures ANOVA in each hemisphere. The ANOVA revealed two left-lateralised effects of COMP in our test window of 0–800ms after onset of the target stimulus, with sentences eliciting more activity than lists (see Figure 6a-b). The first cluster (*p* = 0.0043) extended from ∼220ms to 415ms and had a broad fronto-temporal distribution. The second cluster (*p* = 0.047) occurred slightly later at ∼505ms to 620ms and included additional ventral posterior temporal areas. We did not observe any significant clusters in the right hemisphere.

#### Anterior, then posterior, left fronto-temporal effects of manipulating the syntactic frame

The second round of analyses was specifically aimed at finding effects of our syntactic dependency manipulations unobscured by the list stimuli, and repeated-measures ANOVAs with factors SYN and ARG were therefore run on the sentence stimuli in isolation. This revealed two effects of SYN (see Figure 7a-b); first with more for activity yes/no questions than declarative sentences in inferior frontal and anterior temporal regions (∼220-290ms) and subsequently with more activity for declarative sentences than yes/no questions in a larger cluster also including posterior temporal regions (∼520–635ms).

**Figure 7.**
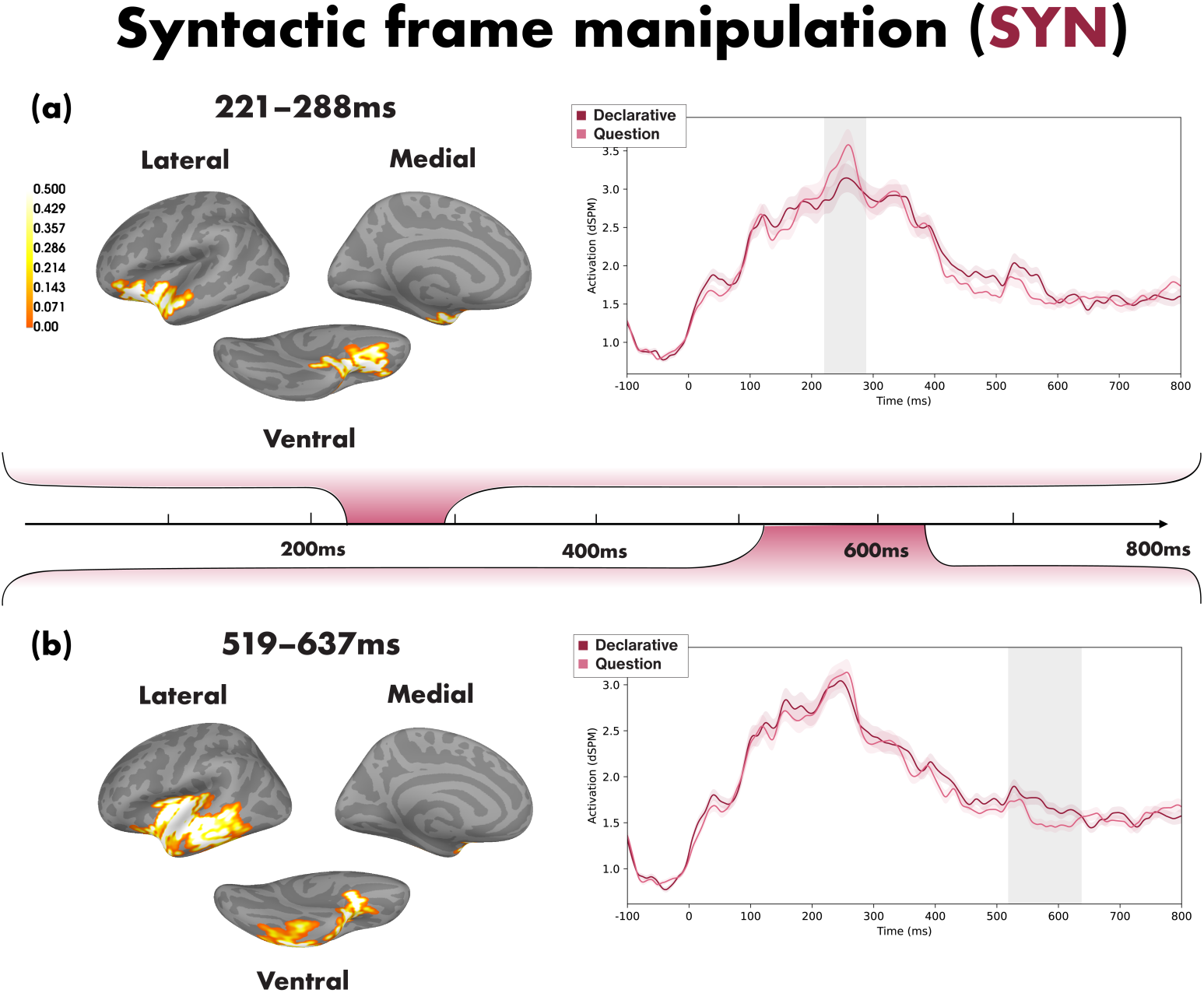
Timeline showing neural effects of manipulating SYN, with the time course for each effect plotting the averaged activity in the identified spatial cluster and the light grey-shading indicating the extent of the significant clusters. (**a)** The first effect of SYN localised in anterior temporal and inferior frontal areas from ∼220–290ms, with yes/no questions eliciting more activity than declarative sentences, while (**b**) the second effect of SYN had a broader temporal distribution from ∼520–635ms with the reverse directionality. A post-hoc analysis on the list stimuli replicated the first SYN effect but not the second one. SYN: syntactic frame; * = p < 0.05 corr.

First, we performed targeted investigations of our syntactic dependency manipulations in each of the clusters identified as instances of the Sentence Superiority Effect. Because the second cluster already included large swaths of the left temporal lobe, we chose to include the entire temporal lobe in this fROI given the general sensitivity of this region to compositional—and especially syntactic—processes. We did not expand the first cluster as it only engaged the very anterior parts of the left temporal lobe. Each fROI yielded a significant effect of SYN, respectively in the early (100–500ms) and late (500–800ms) time windows. In the first fROI, the cluster (*p* = 0.0417) localised in anterior temporal and inferior frontal areas at ∼220-290ms, with yes/no questions eliciting more activity than declarative sentences. The opposite pattern of activation was found for the cluster in the second fROI, with activity increases for declarative sentences over yes/no questions across inferior frontal regions and the majority of the temporal lobe at ∼520–635ms (*p* = 0.0189). To unpack the SYN effects, we performed post-hoc tests following the same analysis pipeline on the list stimuli only. While the early SYN effect replicated for lists (*p* = 0.0133), we saw no such replication for the later SYN effect. This suggests that at least the latter effect must originate in linguistic computations rather than only reflecting swapping the word order.

Our bilateral lobe-by-lobe search in the early (100–500ms) time window did not yield any effects that remained significant after correcting for multiple comparisons, but the same search in the later time window (500–800ms) revealed an effect of SYN at ∼520–635ms (unccor. *p* = 0.0115, corr. *p* = 0.046). Again, the effect did not emerge when performing a post-hoc test on the list stimuli only. Since this cluster had a spatiotemporally similar profile to the later SYN effect observed in the fROI analysis, we take them to reflect the same neural activity and will simply refer to the fROI results going forward. All remaining clusters did not reach the threshold for significance (corrected *p* < 0.05). Our analyses looking for effects across each hemisphere in the entire epoch did not reveal any effects. In sum, our syntactic dependency manipulations yielded an early and a late effect of SYN with different directionality while we did not see any effects of ARG.

## DISCUSSION

In this study, we employed a novel combination of Danish stimuli and Rapid Parallel Visual Presentation (RPVP) to explore the neural signatures of syntactic dependency manipulations when freed from the temporally induced prediction effects and differential condition-specific working memory costs inherent of serial processing. Specifically, we varied the syntactic frame (SYN; declarative sentences, yes/no questions) and the argument structure of intransitive verbs (ARG; unergative, unaccusative, alternating unaccusative). The choice of stimulus delivery technique additionally allowed us to explore, for the first time with minimal two-word sentences and lists (COMP; sentences, lists), the behavioural and neural correlates of the so-called Sentence Superiority Effect. Our main results include behavioural and neural instances of the Sentence Superiority Effect, with facilitated processing of two-word sentences over two-word lists across-the-board, as well as two neural effects arising from our manipulation of syntactic frame, the first with increased activation for yes/no questions and the second with increased activation for declarative sentences. We take the first effect of SYN to reflect basic structure detection and the second effect of SYN to reflect syntactic movement. Below we discuss first the instances of the Sentence Superiority Effect obtained in the present study before turning to the SYN results. Common to all our experimental manipulations, however, is that the neural responses we observe are to some extent determined by the way language was delivered. While seriality is an inherent characteristic of spoken language, we find evidence that the language system is highly flexible and can extract vast amounts of information of varying granularity from input presented in parallel. The fact that we observe such differences even for two-word sentences underscores the language system’s exceptional sensitivity for detecting meaningful grammatical details between minimally distinct stimuli.

### A behavioural and neural Sentence Superiority Effect for minimal structures

This study adds to a growing body of RPVP work, spearheaded by Snell & Grainger (2017), that investigates the Sentence Superiority Effect, a phenomenon where sentences enjoy a processing advantage compared to their unstructured counterparts. We find substantial support for such a Sentence Superiority Effect, notably obtained using shorter stimuli (two-word sentences vs. two-word lists) than those of prior studies. Like prior RPVP literature investigating the behavioural Sentence Superiority Effect, we found reaction times to be shorter and accuracies higher for sentences than for unstructured representations. Such facilitative effects thus suggest that syntactic structure aids comprehension rather than adding cognitive load. Neurally, we first see a broadly distributed instance of the Sentence Superiority Effect in left fronto-temporal regions at around 220–415ms followed by a second effect that included more ventral and posterior temporal areas at around 505-620ms, with sentences eliciting more activity than lists in both cases. Our MEG results thus largely mirror those reported by prior electrophysiological research in terms of timing and, in the case of the comparable MEG studies (Fallon & Pylkkänen, 2024; Flower & Pylkkänen, 2024), directionality. Such directionality was further predicted based on a comprehensive body of MEG research finding increased activity for two-word phrases over two-word lists (see Pylkkänen (2019, 2020) for reviews). Crucially, our behavioural and neural Sentence Superiority Effects are obtained using a simple matching task, which could be performed purely based on perceptual similarity. The fact that we still see a difference between the processing of sentences and lists therefore underscores the language system’s preference for structured representations.

As one of the first studies exploring the neural correlates of the Sentence Superiority Effect, we had no specific hypotheses as to where the effect(s) might localise. It seemed plausible, however, that we would see an effect with a widespread distribution given that our contrast—sentences vs. lists—was a gross one with both syntactic and semantic differences. This notion is supported by prior MEG research (Dufau et al., 2024; Fallon & Pylkkänen, 2024; Flower & Pylkkänen, 2024), who also found broad left-lateral distributions despite using different contrasts to bring about instances of the Sentence Superiority Effect. Our first cluster encompasses typical language-processing regions like the LIFG and the LATL and looks most similar to the neural instance of the Sentence Superiority Effect observed by Flower & Pylkkänen (2024). As regions frequently implicated in, respectively, syntactic processing and conceptual combination (Bemis & Pylkkänen, 2011), it is not unexpected to see an instance of the Sentence Superiority Effect localising here. The second cluster stretches further back and includes more posterior and ventral parts of the temporal lobe, similar to what was observed by Fallon & Pylkkänen (2024). Altogether, our instances of the Sentence Superiority Effect thus encompass a number of quintessential language regions (Fedorenko et al., 2010; Fedorenko & Thompson-Schill, 2014). Notably, however, inferior parietal regions are conspicuous in their absence in our study, even though they participated in the instances of the Sentence Superiority Effect identified by Fallon & Pylkkänen (2024). One possible explanation for this could be our choice of control condition. We chose a two-verb list condition to avoid unintended composition effects arising from the homonymous nature of many Danish nouns and verbs, but it is possible that the verb list-contrast as opposed to the frequently used noun list-contrast from RSVP studies may have washed out significant differences between sentences and lists. This interpretation is supported by the nature of Fallon & Pylkkänen’s (2024) effects; while their first cluster showing a neural Sentence Superiority Effect did not replicate for relative clauses or stimuli with two sources of processing difficulty (agreement errors and semantic role reversals), their second cluster did. As such, it is possible that this latter effect, which had the most extensive involvement of the inferior parietal cortex, was driven by the sheer presence of a verb rather than the detection of structural representations. However, more research using different kinds of contrasts to elicit the Sentence Superiority Effect is needed to clarify this point.

While we find support for the existence of a Sentence Superiority Effect, it also brings to the fore a crucial question: What aspects of sentence processing actually underpin this Sentence Superiority Effect? The ability to bring about instances of the Sentence Superiority Effect by contrasting grammatical sentences with various non-sentence structures—scrambled sentences, reversed sentences, noun lists, verb lists—points to it being of a decidedly structural nature. Even so, the hypothesis space remains vast, and it is therefore a promising avenue for future research to deconstruct. Our syntactic dependency effects, which we will turn to next, shared spatiotemporal loci with our instances of the Sentence Superiority Effect and may therefore provide clues as to the underlying linguistic computations driving our instances of the Sentence Superiority Effect.

### Left fronto-temporal effects of syntactic frame: From structure detection to structure modification

In Danish, declarative sentences and yes/no questions differ exclusively in their word order. This property allowed us to manipulate the syntactic structures of two-word sentences with intransitive verbs like, e.g., the declarative sentence *pigen festede* (“the girl partied”) and the yes/no question *festede pigen* (“did the girl party”), while keeping surface lexico-semantics constant. Current accounts from linguistic theory schematise this difference as a structure with fewer syntactic dependencies for yes/no questions than for declarative sentences and vice versa in terms of merged elements, both arising from the V2 properties of Danish.

Interestingly, the brain is able to detect the minimal difference between declarative sentences and yes/no questions induced by a word order swap, even when the stimuli are presented for a short amount time as done in RPVP. Our MEG data revealed two significant effects of SYN when using the two instances of the Sentence Superiority Effect as fROIs: an early effect with a left inferior frontal-anterior temporal locus at around 220–290ms and a later effect with a broader temporal distribution at around 520–635ms. Post-hoc tests on the list stimuli, which were paired in the order Transitive-Intransitive and Intransitive-Transitive to reflect the syntactic frame manipulation in a structure-free context, replicated the early but not the later effect. Crucially, the two neural effects differed not only in timing and spatial origin but also in directionality. Whereas the first SYN effect saw more activity for yes/no questions than declarative sentences, the opposite pattern was observed for the second SYN effect. Coupled with the spatiotemporal differences, it therefore seems likely that the two clusters reflect different computations.

Before zooming in on the clusters, we wish to highlight that the brain’s detection of the word order swap is noteworthy in itself given the Transposed-Word Effect observed for sentences like *you that read wrong* in prior RPVP research (Flower & Pylkkänen, 2024; Mirault et al., 2018; Pegado et al., 2021; Snell & Grainger, 2019; Wen et al., 2021b). Since Flower & Pylkkänen (2024) found evidence that the brain first detects and then “fixes” such minor word order-errors, it was possible, in the context of our study, that the brain would attempt to “fix” the word order of the less frequently occurring yes/no questions to be like the ubiquitous declarative sentences, but this is not what we see. This, however, is not surprising since Danish readers must necessarily rely on such word-order variation to infer the correct meaning of a sentence. There is thus a stark difference between grammatical word-order variation and ungrammatical word-order errors.

What might each of the two SYN effects be indexing? The lack of prior research tracking the temporal evolution of sentential representations from stimuli presented in parallel, coupled with our novel syntactic frame manipulation juxtaposing declarative sentences with yes/no questions, makes it difficult to directly compare our findings to existing literature, and we therefore draw parallels to other linguistic phenomena with similar spatial and/or temporal profiles in our explanation. Beginning with the first effect, its timing and localisation aligns well with a comprehensive literature finding LATL involvement during basic composition at 200–250ms (Pylkkänen, 2019; 2020) and, in the context of RPVP work using MEG, with the left fronto-temporal instance of the Sentence Superiority Effect observed by Flower & Pylkkänen (2024). Since we controlled bigram probabilities, we can rule out form-based prediction estimates resting on pure word category identification (Noun-Verb and Verb-Noun) associated with these specific lexical items. However, form-based prediction estimates could also rely on basic constituency-detection of licit syntactic structures like Subject-Verb and Verb-Subject (cf. Fallon & Pylkkänen (2024) and Dunagan et al. (2025) for more on the hypothesis of basic constituency-detection). Unfortunately, we cannot ratify such an interpretation given the ability to prepose non-subject constituents like objects and adverbials in V2 languages, making it impossible to dissociate, e.g., Subject-Verb from Noun-Verb structures in Danish. Even so, the directionality of the activity provides further support for this early SYN effect indexing basic constituency-detection: Declarative sentences have syntactic dependencies not found for yes/no questions, thus causing the former to be theoretically further divorced from the base-generated structure than the latter.

What remains curious, however, is that the first effect of SYN arises from two-word sentences and two-verb lists alike; since the latter was intended as a non-combinatory control, effects reflecting linguistic computations should not replicate for the list stimuli. We speculate that this may be an unintended consequence of length-matching the transitive verbs used in the lists with the nouns used in the sentences, which grouped together nouns and transitive verbs (5–7 letters long) in opposition to intransitive verbs (7–10 letters long). In other words, the neural signals may reflect form-based cues to the position of the intransitive verb regardless of whether it occurred in a sentence or a list. More research is needed to disentangle the two potential interpretations from one another.

The second SYN effect localised in the temporal lobe, involving both anterior and posterior regions, as well as in neighbouring inferior frontal regions. This replicates the findings of a substantial body of neuroimaging literature implicating the LPTL and the LIFG in syntactic processing (e.g., Matchin et al., 2017, 2019; Pallier et al., 2011; Zaccarella & Friederici, 2015; Zaccarella et al., 2017; see also Matchin & Hickok (2020) for a recent review). The latency of the effect suggests that it may involve modification of an existing structure, not least because we see evidence of basic constituency-detection already in our early instance of the Sentence Superiority Effect. Such an interpretation is further supported by the fact that declarative sentences, which involve an additional syntactic dependency compared to yes/no questions, saw activity increases in this second SYN effect.

In terms of timing, our SYN effect is somewhat late compared to prior MEG research implicating the LPTL in syntactic processing; for example, Flick & Pylkkänen (2020) and Matar et al. (2021) both report elevated LPTL activation at 200–300ms after target word-onset. Furthermore, they saw increased activation for simpler syntactic structures over more complex ones, thus contrasting with the nature of our SYN effect. However, these differences may be rooted in the different experimental designs. First, Flick & Pylkkänen (2020) as well as Matar and colleagues (2021) both used RSVP for their stimulus presentation, thus allowing incremental structure building as the stimulus unfolds. Our study, in contrast, presented the entire stimulus at once, rendering incremental structure building impossible. It is not unreasonable to presume that at least some syntactic operations will be delayed until a basic syntactic structure has been established, thereby explaining the latency differences. Accounting for the difference in directionality is slightly more complicated but can be done at least in the case of Flick & Pylkkänen’s (2020) study. They contrasted sentences with postnominal predication (*Arei many trails ti wide enough …*) and sentences with postnominal modification (*There are many trails wide enough …*), thus varying the extent of structure required to close the syntactic node containing the adjective, and found more activity for postnominal predication-sentences which allowed for immediate syntactic composition on the adjective. Interestingly, however, these sentences included a syntactic dependency (indicated by the trace *ti*) not found in sentences with postnominal modification, similar to our declarative sentence vs. yes/no question-contrast. Accordingly, it is possible that elevated LPTL activity can be related to some forms of syntactic dependency resolution; however, this remains a question for future studies.

At least one non-syntactic hypothesis regarding the origins of the SYN effects ought to be entertained, namely the actual meanings of the sentences. Given its latency, the second SYN effect is a particularly strong candidate for indexing high-level processing related to interpreting the sentence as either a statement or a question. Here it is again relevant to consider the MEG study by Flick & Pylkkänen (2020). Their stimuli (*Arei many trails ti wide enough …* vs. *There are many trails wide enough …*) likewise differed in both syntactic complexity as well as discourse-level interpretations, and they therefore included a control experiment mimicking the declarative vs. interrogative sentence-contrast of their main experiment while keeping syntactic structure constant. Their LPTL effect did not surface in the control experiment, suggesting that it did not reflect the manipulation of interrogative force. Thus a syntactic interpretation of our SYN effect in the LPTL is also more likely than the semantic alternative.

#### No neural effects of verb argument structure

Intransitive verbs come in different flavours based on the thematic roles they assign to their arguments, which in turn reflects distinct syntactic structures. While the subject of an unergative verb like *party* originates in the subject position, unaccusative and alternating unaccusative verbs like *awake* and *dry* attract the object-generated argument to the subject position. As it turns out, we did not observe any neural effects of ARG. This could be expected as the success rate in eliciting differential neural responses to this manipulation varies substantially across the existing neuroimaging literature (Agnew et al., 2014; Martinez de la Hidalga et al., 2019; Meltzer-Asscher et al., 2013, 2015; Shetreet et al., 2009; Shetreet & Friedmann, 2012; Wang et al., 2021; Zawiszewski et al., 2022). Given the prior RPVP literature, it is furthermore quite possible that the lack of ARG effects stems from a reliance on rather shallow form-related cues to structure during rapid processing. Since unergative, unaccusative, and alternating unaccusative verbs share the same Subject-Verb surface order, a hypothesis relying on basic constituency-detection would not predict any differential processing of the verbs, even if they have different underlying structures (cf. Fallon & Pylkkänen (2024) and Dunagan et al., (2025)). That said, as with any interpretations hinging on a null effect, further investigations of syntactic structure building using RPVP are needed to probe if other superficially identical but underlyingly different stimuli likewise fail to produce distinct variations in neural activity.

## CONCLUSION

What are the neural correlates of syntactic dependencies when dissociated from the temporally induced incremental prediction effects and differential working memory loads across experimental conditions that typically accompany research using serial stimulus delivery techniques? Here we combined properties of Danish syntax with Rapid Parallel Visual Presentation (RPVP) and a simple matching task to address these challenges in a study varying the type (syntactic frame: declarative, yes/no question; verb argument structure: unergative, unaccusative, alternating unaccusative) and number of syntactic dependencies in two-word sentences. Like prior RPVP studies, we found behavioural and electrophysiological “Sentence Superiority Effects”, realised as faster and more accurate responses for sentences than lists controls and left fronto-temporal activity increases (221–417ms; 506–622ms) for sentences. This is the first report of Sentence Superiority Effects for such minimal structures. Our syntactic manipulations induced two left fronto-temporal effects of syntactic frame—first with an anterior locus (221–288ms) and later with a broader temporal distribution (519–637ms)—hypothesized to reflect basic constituency-detection and subsequent syntactic dependency resolution. We did not find effects of our verb argument structure manipulation. The neurobiology of displacement may thus benefit from investigations using RPVP to delineate the impact of covarying factors typically associated with serial stimulus presentation on syntactic processing. Together, our results underscore the swift and automatic nature of the language system that—based on input presented for just a few hundred milliseconds—can generate representations sophisticated enough to elicit differential neural responses to superficially similar but linguistically distinct stimuli.

## ACKNOWLEDGEMENTS

This work was supported by the National Science Foundation award #2335767 (LP) and award G1001 from NYUAD Institute, New York University Abu Dhabi (LP). We thank Alec Marantz and Ailís Cournane for comments on earlier versions of this manuscript and the members of the Neuroscience of Language Lab at NYU for helpful discussion and feedback throughout.

## Box 1.

Example sentences from Danish illustrating some of the constituents that may occupy the first position in main clauses

### Subject in first position

*Hun vandt heldigvis maratonet.* She won luckily marathon-the. ”She luckily won the marathon.”

### Object in first position

*Maratonet vandt hun heldigvis.* Marathon-the won she luckily. “The marathon she luckily won.”

### Adverb in first position

*Heldigvis vandt hun maratonet.* Luckily won she marathon-the.

”Luckily, she won the marathon.”

### Embedded clause in first position

*Selvom hun vågnede for sent, vandt hun heldigvis maratonet*.

Even-though she awoke too late, won she luckily marathon-the. “Even though she woke up too late, she luckily won the marathon.”

## Box 2.

Diagnostics used to classify verbs as unergative, unaccusative, or alternating unaccusative

Verbs were classified as unergative (i) if they employed the auxiliary *have* (“to have”) in the perfect, (ii) if they could occur with a reflexive pronoun in resultative constructions, and (iii) if they could occur in impersonal passive constructions. Additionally, unergative verbs had to fail (iv), (v), and (vi).

(i) *Pigen har festet*. “The girl has partied.”
(ii) *Pigen festede sig selv træt*. “The girl partied herself tired.”
(iii) *Der festes*. “Someone is partying.”
(iv) * *Den festede pige*. “The partied girl.”
(v) * *Pigen festede en fest*. “The girl partied a party.”
(vi) * *Pigen blev festet*. “The girl was partied.”

Verbs were classified as unaccusative (i) if they employed the auxiliary *være* (“to be”) in the perfect, and (iv) if past participle forms could be used as nominal modifiers. Additionally, unaccusative verbs had to fail (ii), (iii), (v), and (vi).

(i) *Pigen er vågnet*. “The girl is awoken.”
(ii) * *Pigen vågnede sig selv frisk*.
*\* Der vågnes*.
*Den vågne pige*.
* *Pigen vågnede en kat*.
* *Pigen blev vågnet*.

Verbs were classified as alternating unaccusative (i) if they could take either auxiliary in the perfect, (iii) if they could occur in impersonal passive constructions, (iv) if past participle forms could be used as nominal modifiers, (v) if they could occur with direct objects, and (vi) if they could undergo passivization. Alternating unaccusative verbs generally did not fare well with reflexive pronouns in resultative constructions as in (ii).

(i) a. *Pigen er tørret*. “The girl is dried.”
b. *Pigen har tørret blusen*. “The girl has dried the shirt.”
(ii) ? *Pigen tørrede sig selv varm*.
(iii) *Der tørres*.
(iv) *Den tørrede pige*.
(v) *Pigen tørrede en bluse*.
(vi) *Pigen blev tørret*.

